# Obsessive Compulsive Disorder and Response Inhibition: Meta-analysis of the Stop-Signal Task

**DOI:** 10.1101/2021.07.16.452538

**Authors:** Kendall Mar, Parker Townes, Petros Pechlivanoglou, Paul Arnold, Russell Schachar

**Affiliations:** Hospital for Sick Children (SickKids), University of Toronto; Matheson Centre for Mental Health Research and Education, Hotchkiss Brain Institute, University of Calgary

**Author notes:** **Author Note**, Kendall Mar is now at the University of Toronto, Department of Psychology. All authors declare that they have no conflicts of interest. The authors wish to thank Quenby Mahood for assistance in the literature search strategy. The manuscript has been pre-registered online prior to peer review through PROSPERO (https://www.crd.york.ac.uk/prospero/display_record.php?RecordID=212770). Correspondence concerning this article should be addressed to Russell Schachar, Hospital for Sick Children (SickKids), University of Toronto, 555 University Ave, Toronto, ON M5G 1X8, Canada.

**Keywords:** Obsessive-Compulsive Disorder, Impulsive Behavior, Stop Signal Task, Response Inhibition, Executive Function, Response Time

## Abstract

This systematic review and meta-analysis updates evidence pertaining to deficient response inhibition in obsessive-compulsive disorder (OCD) as measured by the stop-signal task (SST). We conducted a meta-analysis of the literature to compare response inhibition in patients with OCD and healthy controls, meta-regressions to determine relative influences of age and sex on response inhibition impairment, and a risk of bias assessment for included studies using the Newcastle-Ottawa Scale (NOS). Stop-signal reaction time (SSRT), which estimates the latency of the stopping process deficit, was significantly longer in OCD samples than in controls, reflecting inferior inhibitory control (Raw mean difference = 23.43ms; p = <0.001; 95% CI = [17.42, 29.45]). We did not observe differences in mean reaction time (MRT) in OCD compared to controls (Raw mean difference = 2.51ms; p = 0.755; 95% CI = [−13.27, 18.30]). Age impacted effect size of SSRT, indicating a greater deficit in older patients than younger ones. We did not observe a significant effect of sex on SSRT or MRT scores.

**General Scientific Summary:** Difficulty inhibiting responses is an hypothesized deficit in Obsessive-Compulsive Disorder (OCD). The results of this systematic review and meta-analysis of studies using the Stop Signal Task support the notion of deficient response inhibition in OCD and indicate that older individuals with OCD show greater impairments than younger ones.

## Introduction

OCD is a debilitating and chronic mental illness characterized by obsessions and compulsions (American Psychiatric Association, 2013). The worldwide lifetime prevalence of OCD is approximately 1-4% for adults (Kessler et al., 2012; Ruscio et al., 2010; Samuels & Nestadt, 1997) and for children (Grados et al., 1997; Helbing & Ficca; 2009; Samuels & Nestadt, 1997). Treatment of OCD can include medication or cognitive behavioral therapy. The cause of OCD is poorly understood (Fernandez & Leckman, 2016), but is likely to involve a complex interplay of psychosocial and neurobiological factors including genetic and neural risks (Bandelow et al., 2016; Monzani et al., 2014), which might differ between children and adults (Mattina & Steiner, 2016; Taylor, 2011). Deficient executive control, in particular response inhibition, may mediate the link between underlying risk, the ability to control thought and actions, and OCD symptoms (Robbins et al., 2019).

For example, response inhibition, the ability to stop an action or thought when one’s goals or external circumstances change, is an aspect of executive control that is of considerable interest in OCD (Dupuy et al., 2013). In the absence of adequate response inhibition, thoughts and actions are more difficult to control and less likely to be stopped so that they can be replaced with more adaptive thoughts or actions. Response inhibition has been likened to the ability to slam on the car’s brakes when someone steps into the road or to check a swing in baseball.

The stop-signal task (SST) is one of the most common laboratory paradigms for studying inhibition of a speeded motor response. The SST involves presentation of go and stop signals. On ‘go’ trials, the participant is presented with a ‘go’ stimulus on a computer screen and is instructed to respond as quickly and as accurately as possible with either a right-hand or left-hand response. On a random subset of trials (typically about 25%), the ‘go’ stimulus is followed by a ‘stop’ signal (usually a tone presented through headphones). The participant is instructed to stop their response when they hear the stop signal. The delay between presentation of the ‘go’ and presentation of the ‘stop’ signal is dynamically adjusted depending on participant’s performance. If a response is successfully stopped, the delay increases, making it more difficult to stop on subsequent ‘stop’ trials. If a response is not stopped, the delay decreases making it easier to stop on subsequent trials. Tracking proceeds to the delay at which the participant can stop half the time called the stop signal delay (SSD). Stop-signal reaction time (SSRT) is the latency in milliseconds of the otherwise unobserved stopping process (Logan & Cowan, 1984; Logan et al., 1984). Longer SSRT indicates deficient response inhibition.

Although there have been several hundred studies using the SST in disorders such as attention-deficit hyperactivity disorder (ADHD), there is far less research in OCD (Lipszyc & Schachar, 2010). The Lipszyc and Schachar (2010) review could find only four relevant studies and meta-analysis which revealed a deficit in response inhibition of medium effect size in OCD patients when compared to healthy controls. Abramovitch and colleagues (2013) also conducted a review of the literature concerning response inhibition in individuals with OCD compared to healthy controls and found a significant difference in effect size. Their study however, included a multitude of tasks to measure response inhibition, but had incomplete coverage of the literature, and failed to review the most important dependent variables from the SST (Abramovitch, et al., 2013). Eng and colleagues (2015) also completed a review investigating differences in structural grey matter activation, where they did not find a significant difference in activations during response inhibition tasks in individuals with OCD compared to controls. This review only included one study that utilized the SST.

The present systematic review and meta-analysis aims to update what is known about response inhibition in OCD as measured in the SST, as well as determine if there is an association with age and response inhibition impairment. Although SSRT is the main outcome of the SST, the task has been used to measure speed (MRT) and variability (standard deviation of reaction time, RTSD) as indicators of arousal or attention. Differences in MRT have been used to qualify the interpretation of longer SSRT (Alderson et al., 2008). Therefore, we include MRT and RTSD as outcomes in this meta-analysis. Previous meta-analytic studies investigating the SST have identified response inhibition deficits in substance use and addiction (Smith et al., 2014) and ADHD (Alderson et al., 2007; Huizenga et al., 2008) indicated by significantly longer SSRT relative to controls. Therefore, in the present study we hypothesize that individuals with a diagnosis of OCD would have a significant deficit in response inhibition, indicated by a longer SSRT, when compared to age and sex matched healthy controls.

## Methods

We performed a systematic review of the literature in accordance with the Preferred Reporting Items for Systematic Reviews and Meta-Analysis (PRISMA) guidelines (Moher et al., 2009; Liberati et al., 2009), See online Supplement, S4, for full PRISMA checklist. The study protocol was registered with PROSPERO (No. 212770).

The systematic review was limited to case-control studies that fulfilled the following criteria: (a) used the SST task; (b) included a healthy comparison group; (c) reported results from at least one of the SST variables of interest (SSRT, MRT and RTSD) for effect size calculations (i.e. means and standard deviations or F and t values); (d) involved OCD cases that were defined by DSM or ICD classification; (e) published between April 1960 and November 2020; (f) published in a peer-reviewed journal and; (g) had a full-text article available. Studies that included a modified version of the SST were permitted if the paradigms were administered like the typical SST and the outcome measures of interest were discussed. Studies with a mean age difference of three years or more between participant and control groups, those that failed to explicitly state that participants were not on stimulant medication due to its potential to effect on an individual’s reaction time (Pliszka et al., 2007; Tannock et al., 2000), and those that provided feedback to participants, due to the suspected effect of reward on response inhibition (Sinopoli et al., 2011; Slusarek et al., 2001) were excluded from the study.

Studies were identified by a systematic and comprehensive literature search of the Embase OVID, MEDLINE, PsychINFO and Web of Science electronic databases. The search strategy utilized to identify potential studies was created in collaboration with a librarian. See online Supplement, S1, for the full search strategy. Studies that were included in previous meta-analyses of the SST, response inhibition and OCD were considered for inclusion into the present meta-analysis. Reference lists of relevant articles were also reviewed to further exhaust the literature.

Retrieved literature from the search underwent initial eligibility screening of titles and abstracts by two independent raters (PT & KM) who were blinded to each other’s decisions. Following title and abstract screening, the full-text of all candidate articles were obtained and assessed independently for inclusion by two blinded independent raters (PT & KM). A third rater (RS) resolved discrepancies between reviewers.

Reviewers (PT & KM) independently extracted data from eligible studies using a form based on the Cochrane Handbook Chapter 5 (Li et al., 2020). Information extracted from each study included study design and location, demographic characteristics of the cohort, mean SSRT and standard deviation, MRT and standard deviation and RTSD scores. Discrepancies were resolved through consensus between the two reviewers. A third rater (RS) resolved disagreements between reviewers. The corresponding author was contacted directly for results where data was not reported in the published paper.

We used the Newcastle-Ottawa quality assessment scale (NOS) for case-control studies to assess the risk of bias for each included article (Wells et al., 2019). The NOS was selected for its applicability to non-randomized case-control studies (Zeng et al., 2015) and good validity and reliability (Wells et al., 2019). Within the comparability of cases and controls section of the NOS, reviewers are given the freedom to select two comparability factors that are most relevant to their review. In the present study, we selected age and sex as our comparability factors. Risk of bias scores did not influence article selection or inclusion for the current review.

Reviewers (PT & KM) blind to each other’s decisions conducted the NOS for each of the studies using an online form that was created based on the NOS coding manual for case-control studies (Wells et al., 2019). A single star was allotted for each criterion fulfilled by the study where a greater number of stars allotted represents a lower risk of bias score for the study, up to a maximum score of nine stars. Discrepancies between reviewers were resolved through consensus.

SSRT was the primary outcome of interest in the present meta-analysis, with RTSD and MRT as the secondary outcomes of interest. A weighted random-effects meta-analysis model was performed as the primary outcome measure in SSRT, SDRT and MRT scores between OCD participants and control groups. We report our results as a raw mean difference, using restricted maximum-likelihood as the model estimator to allow for an intuitively meaningful interpretation of our results, reported in seconds. Restricted maximum-likelihood was selected as the model estimator of effect size for its applicability to mixed-models and a less biased estimation of variance components (Harville, 1974; Patterson & Thompson, 1971). We also report our results in standardized mean difference in the supplemental information, using Hedge’s G as the model estimator to allow for greater generalizability of our results (Takeshima et al., 2014). A large positive effect size indicated greater impairment in the OCD group than in controls. A mixed-effects regression analysis was performed to determine the relative impact of age and sex as moderator variables for the model estimator of effect size between groups while comparing our outcome variables of interest (Fox & Weisber, 2018; R Core Team, 2019; Viechtbauer, 2010). The raw mean difference in our age analysis identified the level of deficit experienced between mean ages of each study cohort, per year. Assumptions of normality were made for distributions of data in primary and secondary analyses. Analyses were conducted using jamovi version 1.2 (The jamovi project, 2020) with the *Conducting Meta-Analyses in R* with the *metafor package* addon (Lakens, 2017; R Core Team, 2019; Viechtbauer, 2010).

We employed the Q statistic to assess whether heterogeneity among studies is higher than expected by chance (Cochrane, 1954). A statistically significant Q value (p < 0.05) indicates that significant excess between-study variation exists. We also used the I^2^ statistic as an indicator for the proportion of total variance among study estimates due to heterogeneity (Higgins & Thompson, 2002). The I^2^ statistic is a transformation of the Q statistic that depends on the degrees of freedom of the sample and is reported as a percentage of the total variability where percentages of 25%, 50% and 75% represent low, medium and high heterogeneity, respectively (Huedo-Medina et al., 2006).

A funnel plot of standard error against effect size for both the raw mean difference and standardized mean difference were generated to visually inspect potential publication bias.

Publication bias is indicated in a funnel plot through asymmetry in the distribution of study results across a plane.

## Results

We identified 394 potentially relevant articles through our database and reference list search. See online Supplement, S1, for the full search strategy of all four databases. A total of 21 studies met our criteria and were included in the systematic review and meta-analysis (Fig. 1 PRISMA Flow Diagram). See online Supplement, S2, for references of studies included in meta-analyses. A third party (RS) resolved six discrepancies between reviewers (PT & KM) during full-text screening.

**Figure 1.**
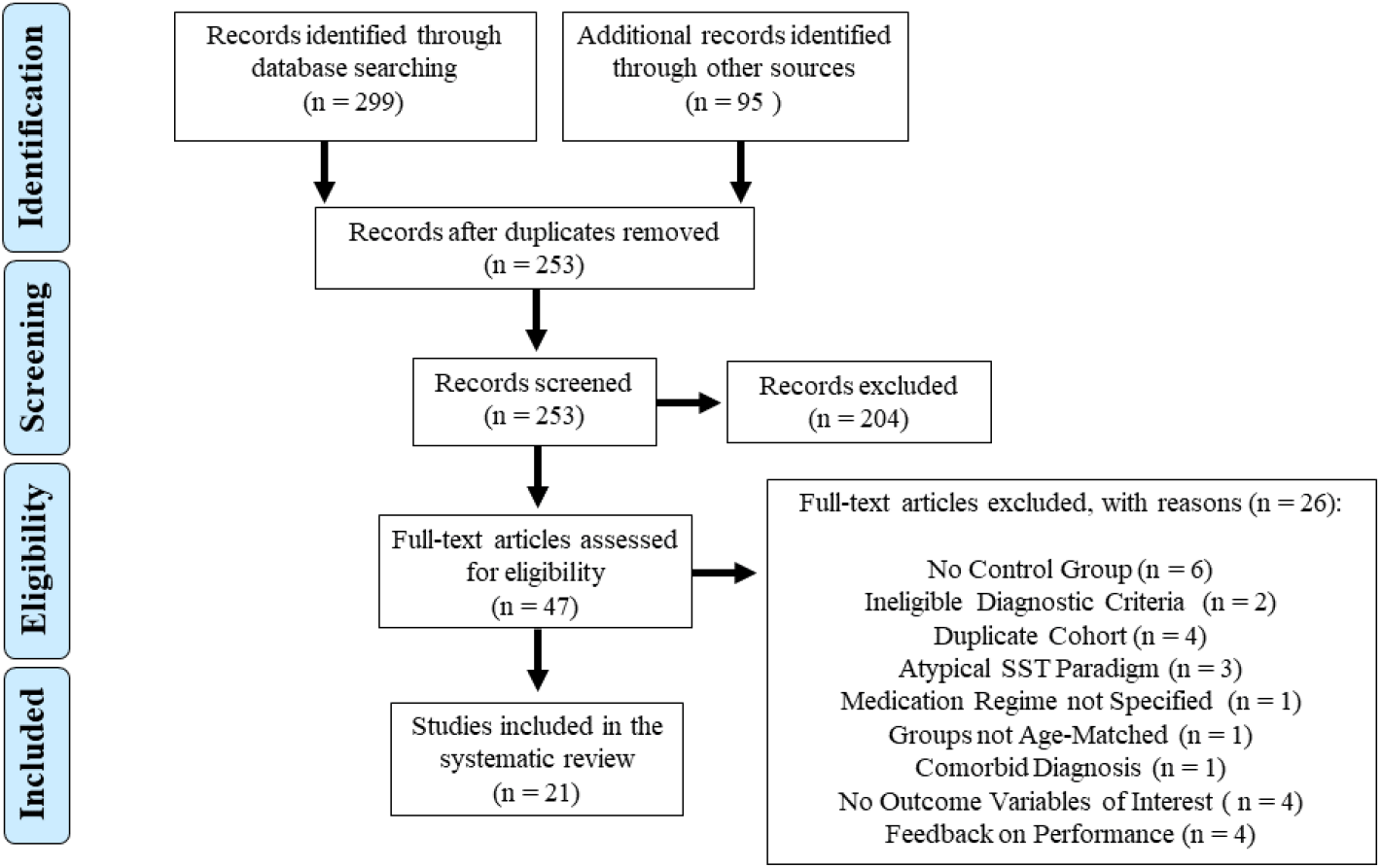
PRISMA Flow Diagram on Article Selection. *Note*. SST = Stop-Signal Task.

Table 1 lists the 21 case-control articles included in this review. The included articles involved a total of 781 OCD and 837 control participants. The average age was 25.77 years in patients with OCD and 24.11 years in healthy controls. OCD participants included a total of 399 females (51.08% of cohort), whereas healthy controls included 412 females (49.22% of cohort). One study did not publish the required data for our analysis, however, the corresponding author provided the necessary information upon request (Heinzel et al., 2018). In two studies (Fan et al., 2016 & Lei et al., 2017), authors included two OCD groups to examine theoretically important research questions and were therefore coded as two separate OCD cohorts.

**Table 1.** Sociodemographic Characteristics of the studies included in the meta-analysis.

| Author<br>(year) | Country | Total | OCD | HC | OCD<br>Age |  | OCD<br>Female |  | HC<br>Age |  | HC<br>Female |  |
| --- | --- | --- | --- | --- | --- | --- | --- | --- | --- | --- | --- | --- |
|  |  | N | N | N | M | SD | N | % | M | SD | N | % |
| Bersani<br>(2013) | Italy | 46 | 25 | 21 | 36.44 | 12.29 | 19 | 76 | 36.85 | 13.27 | 11 | 52.4 |
| Boisseau<br>(2012) | USA | 32 | 19 | 21 | 22.32 | 4.24 | 19 | 100 | 24.24 | 3.47 | 21 | 100 |
| Chamberlain<br>(2006) | UK | 40 | 20 | 20 | 35.3 | 14.0 | 16 | 80 | 34 | 8.1 | 16 | 80 |
| Chamberlain<br>(2007) | UK | 40 | 20 | 20 | 32.1 | 11.9 | 16 | 80 | 33.1 | 10.5 | 13 | 65 |
| De Wit<br>(2012) | Netherlands | 78 | 41 | 37 | 38.6 | 9.8 | 20 | 49 | 39.7 | 11.6 | 19 | 51 |
| Fan (2016)<br>AO | China | 104 | 42 | 62 | 20.36 | 4.68 | 18 | 42.9 | 20.31 | 1.96 | 24 | 38.7 |
| Fan (2016)<br>RO | China | 117 | 55 | 62 | 22.37 | 8.77 | 21 | 38.9 | 20.31 | 1.96 | 24 | 38.7 |
| Frydman<br>(2020) | Brazil | 34 | 17 | 17 | 35.88 | 13.13 | 3 | 17.6 | 35.29 | 10.84 | 3 | 17.6 |
| Gooskens<br>(2018) | Netherlands,<br>Germany<br>and the UK | 83 | 23 | 61 | 10.80 | 1.44 | 12 | 52.2 | 10.76 | 1.2 | 28 | 45.9 |
| Hampshire<br>(2019) | UK | 40 | 20 | 20 | 37.6 | 14.6 | 3 | 15 | 36.3 | 8.3 | 5 | 25 |
| Heinzel<br>(2018) | Germany | 11 | 51 | 49 | 33.00 | 9.73 | 25 | 49 | 30.92 | 7.31 | 29 | 59.2 |
| Kang (2013) | Korea | 36 | 18 | 18 | 24.9 | 5.9 | 6 | 33.3 | 24.7 | 2.7 | 6 | 33.3 |
| Lei (2015) | China | 84 | 42 | 42 | 21.67 | 4.76 | 19 | 45.2 | 22.70 | 3.57 | 18 | 42.8 |
| Lei (2017)<br>early-onset<br>OCD | China | 114 | 63 | 51 | 20.68 | 3.54 | 40 | 63.5 | 21.16 | 4.23 | 33 | 64.7 |
| Lei (2017)<br>late-onset<br>OCD | China | 84 | 33 | 51 | 24.64 | 5.56 | 23 | 69.7 | 21.16 | 4.23 | 33 | 64.7 |
| Negreiros<br>(2020) | Not reported | 166 | 87 | 79 | 13.38 | 2.71 | 61 | 70 | 12.18 | 3.33 | 44 | 56 |
| Ornstein<br>(2010) | Canada | 38 | 14 | 24 | 12.92 | 1.89 | 10 | 71.4 | 12.79 | 2.5 | 13 | 54.2 |
| Penades<br>(2007) | Not reported | 52 | 27 | 25 | 33.7 | 10.5 | 9 | 33.3 | 30.9 | 3.3 | 8 | 32 |
| Sohn (2014) | Korea | 156 | 80 | 76 | 27.8 | 8.1 | 21 | 26.2 | 25.5 | 4.1 | 26 | 34.2 |
| Sunol (2019) | Spain | 37 | 18 | 19 | 46.67 | 9.57 | 8 | 44.4 | 46 | 8.85 | 9 | 47.4 |
| Thorsen<br>(2020) | Norway | 57 | 31 | 26 | 30.19 | 9.21 | 19 | 61 | 31 | 10.73 | 18 | 64 |
| Woolley<br>(2008) | UK | 19 | 10 | 9 | 14.3 | 1.7 | 0 | 0 | 14.5 | 1.1 | 0 | 0 |
| Yu (2019) | China | 52 | 25 | 27 | 24.70 | 7.93 | 11 | 44 | 26.97 | 7.63 | 11 | 40.7 |

*Note.* N = 21 included studies, 23 included cohorts. Two studies (Fan et al., 2016 & Lei et al., 2017) had two distinct OCD groups and were coded as two separate cohorts for our analysis of the stop-signal task.

Raw mean difference results of individual studies are presented in Figure 2(a) for SSRT (n = 21) and Figure 2(b) for MRT analyses (n = 19). The mean difference was significant in OCD and control group SSRT scores (Raw mean difference = 23.4 ms; p = <0.001; 95% CI [17.42, 29.45]). However, the mean difference for OCD patients and control MRT scores saw little variation (Raw mean difference = 2.51ms; p = 0.755; 95% CI [−13.27, 18.30]). Only one study reported RTSD precluding the performance of a meta-analysis. See online Supplement, S6 for the forest plots and S7 for funnel plots in standardized mean difference for both the SSRT and MRT. Our results saw little variation in significance between raw mean difference and standardized mean difference evaluations.

**Figure 2a.**
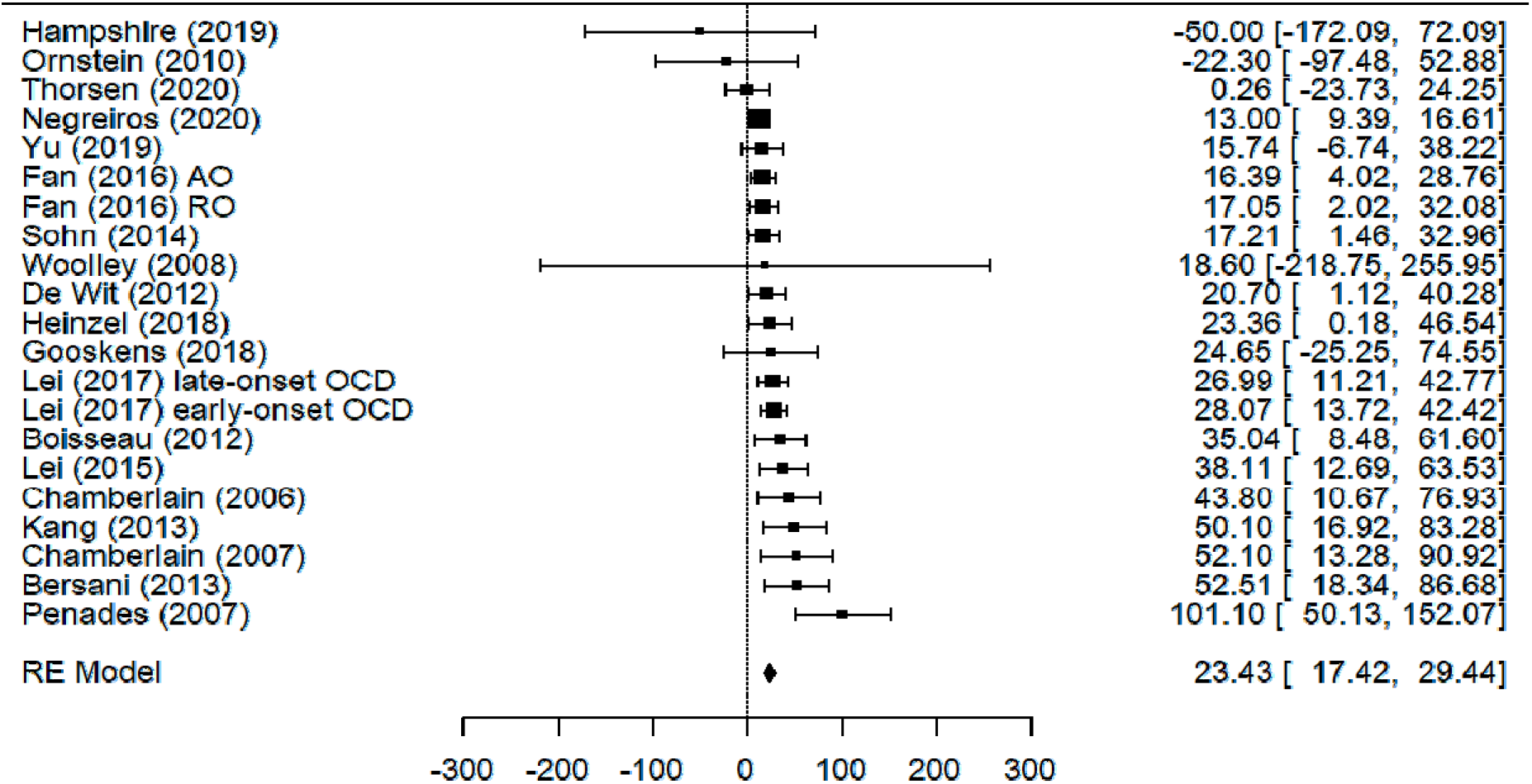
Forest Plot of Included Literature Concerning SSRT Scores Between OCD Patients and Healthy Controls. *Note*. This figure demonstrates the weighted random effects model of the raw mean difference scores of SSRT (n= 21). RE = random effects, as estimated through restricted maximum likelihood. AO = autogenous obsessions. RO = reactive obsessions.

**Figure 2b.**
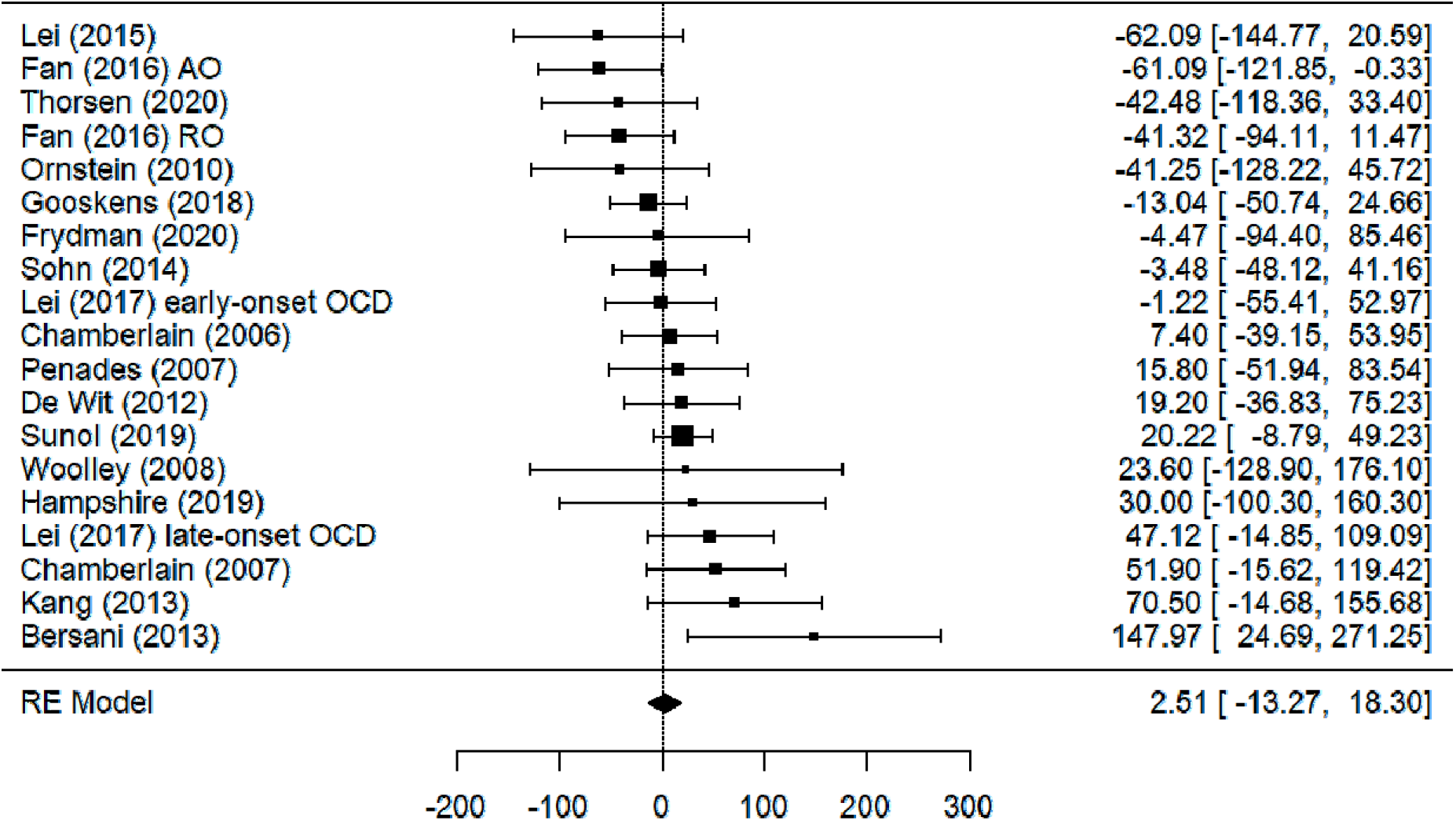
Forest Plot of Included Literature Concerning MRT Scores Between OCD Patients and Healthy Controls. *Note*. This figure demonstrates the weighted random effects model of the raw mean difference scores for studies that included MRT (n= 19). RE = random effects, as estimated through restricted maximum likelihood. AO = autogenous obsessions. RO = reactive obsessions.

Heterogeneity of mean differences between studies was found to be significant for SSRT (Q = 41.62; p = 0.003) and nonsignificant for MRT (Q = 26.33; p = 0.092). The proportion of excess variation between the studies was medium for the SSRT analysis (I^2^ = 41.00%) and low for the MRT analysis (I^2^ = 21.01%). These results indicated a need to examine moderators that could be influencing cohort differences in SSRT mean scores.

Mean age was related to effect sizes for SSRT scores (Raw mean difference = 0.73ms; p = 0.01; 95% CI [0.17, 1.23]) and for MRT scores (Raw mean difference = 1.37ms; p = 0.017; 95% CI [0.24, 2.49]). These results suggest a higher mean age corresponds to a greater deficit in response inhibition and mean response time by 0.73ms and 1.37ms per year, respectively. Correlations between sex (% female) and effect size of SSRT were nonsignificant (Raw mean difference = 0.121ms; p = 0.497; 95% CI [−0.23, 0.47]) as were correlations between sex (% female) and effect size of MRT scores (Raw mean difference = 0.34ms; p = 0.37; 95% CI [−0.48, 1.28]). Taken together, these results indicate that greater mean age but not sex are associated with a more severe impairment in response inhibition.

The risk of bias funnel plot assessment across studies for the random-effects meta-analysis of SSRT scores revealed significant asymmetry (Z = 3.66; *p* < 0.001) (Fig. S5a). Because funnel plot asymmetry analyses typically have low power, however, we cannot conclude the significant result is due to bias alone (Sterne et al., 2011). We suggest this significance may be due to two studies that fall outside the funnel plot (Negreios et al., 2020; Penades et al., 2007) (Figure S5a). After conducting an additional supplemental risk of bias assessment that excluded both studies from the analysis, a nonsignificant heterogeneity of mean differences (Q = 20.868, *p* = 0.286), low proportion of excess variation (I^2^ = 0%) and nonsignificant funnel plot asymmetry (Z = 0.735, *p* = 0.462) were revealed, while maintaining a significant effect size (Raw mean difference = 23.1ms; p =<0.001; 95% CI [18.06, 28.19]) (Fig. S5c). Risk of bias for the MRT analysis demonstrated no evidence of publication bias (Z = 0.77; *p* = 0.439) (Fig. S5b). Upon evaluation of risk of bias between studies using the NOS, all included articles received no less than four out of nine stars, where 157 out of a maximum of 198 stars were awarded (Fig. S3).

## Discussion

The present systematic review and meta-analysis investigated response inhibition as measured in a widely used laboratory paradigm, the stop-signal task, with a focus on the dependent variables of latency of inhibition (SSRT), overall speed (MRT) and variability (RTSD). We sought to update the reliability of evidence for the existence of an inhibition deficit in those with OCD and examine the possibility that age and sex are associated with this deficit.

By comparison to previous reviews, we were able to identify far more studies that investigated response inhibition in OCD using the SST (Abramovitch et al., 2013; Eng et al., 2015; Lipszyc and Schachar, 2010). The literature confirmed the presence of a deficit in response inhibition in OCD. We found no significant differences between individuals with OCD and healthy controls in mean reaction time, suggesting that overall slowing of all responses is not a satisfactory explanation for the longer response inhibition observed in OCD (Alderson et al., 2008). This result suggests OCD participants do not assume a more cautious response set on the SST. By exemplifying similar MRT scores and significantly slower SSRT scores in individuals with OCD compared to healthy controls, our results suggest poorer response inhibition in OCD, thereby supporting the hypothesis of our study. The supplemental results reported in standardized mean difference were comparable to those reported in raw mean difference. We can therefore infer that response inhibition deficits are associated with a diagnosis of OCD.

Inhibition of a speeded motor response is only one of several varieties of inhibition (cf. distractor inhibition, inhibition primed responses) and inhibition is only one form of executive control that humans deploy in their search for self-regulation and goal attainment (e.g. working memory, switching). Although the SST involves stopping a motor response, there is evidence to suggest that the neural circuitry involved may be the same as that involved in stopping of thought as measured in a SST analogue called the Think/No-think task (Guo et al., 2018). Stopping an action in the SST and a thought in the Think/No-think task activates a pathway involving the right inferior frontal gyrus and the basal ganglia (Aron et al., 2014; Chevrier et al., 2007). Common neural activation in the SST and the Think/No-think tasks suggests that the act of intentional control manifested in the SST may be a good proxy for control of thought as well as of action. However, the SST is easier to implement, making it advantageous for study of inhibition in health and disease.

The results of the current review suggest that further study of SST in OCD using imaging techniques could help determine if the circuits involved in executive control differ from those involved in typical development and other psychopathologies, and to investigate the effects of interventions, including drug and non-drug treatments, on performance (Thorsen et al., 2020; Chamberlain et al., 2007). The SST can be easily manipulated to determine the effect of attention, such as variations in emotional content of stimuli, priming, and preparation (Chikazoe et al., 2009) and could therefore be extended to study additional aspects of cognition, such as perception, memory and ultimately to thinking which is, due to its widely distributed nature, likely the most difficult process to study.

Response inhibition deficits are not unique to OCD (Lipszyc & Schachar, 2010; van Velzen et al., 2014). Based on this meta-analysis alone, we cannot draw conclusions about the mechanism of the association between OCD and response inhibition. Response inhibition could be a manifestation of various underlying genetic or neural factors (pleiotropic) or might be an epiphenomenon of OCD as with other neuropsychological disorders. Studies of heritability and co-heritability of response inhibition and OCD would be necessary to address this question. Given the effect size observed in the current meta-analysis, the SST cannot be used as a diagnostic proxy for OCD. It is possible that combinations of executive control measures might have greater accuracy in the detection of OCD than any single measure as has proved to be the case with ADHD (Solanto et al., 2001).

Age was a significant predictor of the magnitude of deficit in SSRT for individuals diagnosed with OCD; older individuals with OCD had greater deficit compared to healthy controls. MRT was also found to be significantly slower with age in the OCD cohort compared to healthy controls. A single study included in the present analysis stratified their OCD cohort into early and late onset and did not observe significant differences in performance between the groups, suggesting that age of symptom onset may not be a marker for response inhibition impairment in adults (Lei et al., 2017). We were unable to conduct an analysis of age of onset as a moderator variable for response inhibition due to lack of reporting across included studies. However, an age-related decline in mean reaction time has previously been observed on the SST in healthy individuals (Hsieh & Lin, 2017). Therefore, while an overall decrease in performance on the SST is observed with age in the current study, individuals with OCD appear to experience a greater age effect compared to healthy controls on both SSRT and MRT scores. Direct comparisons between healthy controls and those with OCD across ages would be useful in clarifying these results. Sex did not predict the magnitude of the effect in OCD even though there is some evidence that males and females may differ in SSRT (Dupuis et al., 2019) and previous studies suggest potential sex-dimorphic activation patterns on imaging of motor inhibition tasks (Rubia 2013). There may be a gap in the literature on sex differences in response inhibition deficits in those diagnosed with OCD. Due to lack of stratified data for male and female OCD and healthy control cohorts, we were unable to investigate whether an interaction existed between age and sex. Given the significant age effects observed in the present paper, this would be an interesting direction for future research.

The NOS risk of bias tool revealed a low risk of bias in study level outcomes for the studies included in this review. Publication bias was found to be insignificant for the MRT analysis and significant for the SSRT, as measured through mean difference analysis in the regression test for asymmetry. The SSRT asymmetry data became insignificant upon removing two studies that fell outside the funnel plot, while maintaining a significant effect size in the primary analysis (Negreios et a., 2020; Penades et al., 2007). One of the outliers, a study by Penades and colleagues (2007), did not vary the order of presentation in the battery of cognitive tests, which we suspect could have contributed to the study’s relatively large significant mean difference. The second outlier by Negreiros and colleagues (2020) may have contributed to funnel plot asymmetry due to the study’s small standard error that could have been caused by the recruitment of a very narrow subset of youth OCD patients from an OCD-specialty program at a tertiary care hospital. Overall, the low risk of bias using the NOS and the justification for the asymmetrical funnel plot gives us confidence in the validity and reliability of our results.

Although the inclusion criteria for this meta-analysis were stringent, several differences between studies have the potential to introduce bias. One important consideration is the potential for pharmacological drug effects on response inhibition. Apart from stimulants, we allowed for the inclusion of studies with OCD participants on other medications. Most OCD cohorts were composed of both medicated and unmedicated subjects. Evidence suggests SSRIs may increase specific regional metabolic rates (Gerdelat-Mas et al., 2005) which would be expected to improve performance on cognitive tasks. Of interest, one study found that the SSRI escitalopram improved the ability to inhibit responses in stop-signal trails in healthy participants (Skandali et al., 2018). Comparisons between medicated and non-medicated OCD cohorts have found nonsignificant differences in SSRT (Kalanthroff et al., 2017) and SSD (Penades et al., 2007). We were unable to conduct additional analyses to investigate the effects of medication on response inhibition in the current paper due to a lack of reporting. This would be a valuable area for future investigation.

Other potential sources of bias between studies include variation in testing environments that were not explicitly disclosed. Although one might presume that they were all laboratory studies, results could be influenced by the amount of supervision that participants received. Some but not all studies reported providing practice on the task and there were several differences among the versions of the administered stop-signal tasks. However, previous studies did not find evidence that variability in the administration of the SST affected results (Dimoska & Johnstone, 2008; Hiraoka et al., 2018).

Given our findings of deficient response inhibition in individuals with OCD, future directions for research could examine response inhibition deficits as a biomarker for OCD. Currently a large gap in the literature exists for longitudinal evaluations of response inhibition in individuals diagnosed with OCD, where two studies exist on the matter and hold a relatively short follow-up interval (Bannon et al., 2006; Thorsen et al., 2020). A more thorough longitudinal investigation of response inhibition is warranted to evaluate the prognosis of individual deficits and may help to answer fundamental questions about development and cognitive decline. Obsessive-Compulsive Disorder has high rates of comorbidity with other psychological disorders, including ADHD, which could impact performance on measures of executive function (Arnold et al., 2005). In the present analysis, comorbidity was not consistently reported in the literature, so we were unable to control for it. However, this may be an interesting direction for future investigation. Lastly, an analysis of reaction time following successful or unsuccessful trials on the SST may also provide insight into how error processing differs in OCD compared to healthy individuals, where individuals with OCD could potentially have a slower reaction time following errors, as mentioned by the error detection theory of OCD (Pitman, 1987). A recent meta-analysis that included a number of inhibitory control tasks found more inhibitory control errors in OCD relative to healthy controls (Norman et al., 2019), however, a more in depth analysis of the relationship between reaction time and error detection in OCD is warranted.

## Conclusions

This systematic review and meta-analysis reveals that people with a diagnosis of OCD have impaired inhibitory control compared to healthy controls. We did not observe a deficit in the mean reaction time in OCD when compared to healthy controls. In addition, results suggest that age has a meaningful, positively correlated influence on the magnitude of deficiencies in response inhibition. By contrast, the role of sex was unclear. Overall, these results indicate that inhibition is a suitable direction for future research into biomarkers for the disorder.

## References

Abramovitch, A., Abramowitz, J. S., & Mittelman, A. (2013). The neuropsychology of adult obsessive-compulsive disorder: a meta-analysis. Clinical psychology review, 33(8), 1163–1171. https://doi.org/10.1016/j.cpr.2013.09.004

Alderson, R. M., Rapport, M. D., & Kofler, M. J. (2007). Attention-deficit/hyperactivity disorder and behavioral inhibition: a meta-analytic review of the stop-signal paradigm. Journal of abnormal child psychology, 35(5), 745–758. https://doi.org/10.1007/s10802-007-9131-6

Alderson, R. M., Rapport, M. D., Sarver, D. E., & Kofler, M. J. (2008). ADHD and Behavioral Inhibition: A Re-examination of the Stop-signal Task. Journal of Abnormal Child Psychology, 36(7), 989–998. https://doi.org/10.1007/s10802-008-9230-z

American Psychiatric Association. Diagnostic and Statistical Manual of Mental Disorders, Fifth Edition. Washington, DC: American Psychiatric Association; 2013.

Arnold, P. D., Ickowicz, A., Chen, S., & Schachar, R. (2005). Attention-Deficit Hyperactivity Disorder with and without Obsessive—Compulsive Behaviours: Clinical Characteristics, Cognitive Assessment, and Risk Factors. The Canadian Journal of Psychiatry, 50(1), 59–66. https://doi.org/10.1177/070674370505000111

Aron, A. R., Robbins, T. W., & Poldrack, R. A. (2014). Inhibition and the right inferior frontal cortex: one decade on. Trends in cognitive sciences, 18(4), 177–185. https://doi.org/10.1016/j.tics.2013.12.003

Bandelow, B., Baldwin, D., Abelli, M., Altamura, C., Dell’Osso, B., Domschke, K., Fineberg, N. A., Grünblatt, E., Jarema, M., Maron, E., Nutt, D., Pini, S., Vaghi, M. M., Wichniak, A., Zai, G., & Riederer, P. (2016). Biological markers for anxiety disorders, OCD and PTSD - a consensus statement. Part I: Neuroimaging and genetics. The world journal of biological psychiatry : the official journal of the World Federation of Societies of Biological Psychiatry, 17(5), 321–365. https://doi.org/10.1080/15622975.2016.1181783

Bannon, S., Gonsalvez, C. J., Croft, R. J., & Boyce, P. M. (2006). Executive functions in obsessive-compulsive disorder: state or trait deficits?. The Australian and New Zealand journal of psychiatry, 40(11-12), 1031–1038. https://doi.org10.1080/j.1440-1614.2006.01928.x

Chevrier, A. D., Noseworthy, M. D., & Schachar, R. (2007). Dissociation of response inhibition and performance monitoring in the stop signal task using event-related fMRI. Human brain mapping, 28(12), 1347–1358. https://doi.org/10.1002/hbm.20355

Chikazoe, J., Jimura, K., Hirose, S., Yamashita, K., Miyashita, Y., & Konishi, S. (2009). Preparation to Inhibit a Response Complements Response Inhibition during Performance of a Stop-Signal Task. The Journal of Neuroscience, 29(50), 15870–15877. https://doi.org/10.1523/JNEUROSCI.3645-09.2009

Crosbie, J., Arnold, P., Paterson, A., Swanson, J., Dupuis, A., Li, X., Shan, J., Goodale, T., Tam, C., Strug, L. J., & Schachar, R. J. (2013). Response inhibition and ADHD traits: correlates and heritability in a community sample. Journal of abnormal child psychology, 41(3), 497–507. https://doi.org/10.1007/s10802-012-9693-9

Dimoska, A., & Johnstone, S. J. (2008). Effects of varying stop-signal probability on ERPs in the stop-signal task: do they reflect variations in inhibitory processing or simply novelty effects?. Biological psychology, 77(3), 324–336. https://doi.org/10.1016/j.biopsycho.2007.11.005

Dupuis, A., Indralingam, M., Chevrier, A., Crosbie, J., Arnold, P., Burton, C. L., & Schachar, R. (2019). Response Time Adjustment in the Stop Signal Task: Development in Children and Adolescents. Child development, 90(2), e263–e272. https://doi.org/10.1111/cdev.13062

Dupuy, M., Rouillon, F., & Bungener, C. (2013). Place de l’inhibition dans le trouble obsessionnel-compulsif [The role of inhibition in obsessional-compulsive disorders]. L’Encephale, 39(1), 44–50. https://doi.org/10.1016/j.encep.2012.06.016

Eng, G. K., Sim, K., & Chen, S. H. (2015). Meta-analytic investigations of structural grey matter, executive domain-related functional activations, and white matter diffusivity in obsessive compulsive disorder: an integrative review. Neuroscience and biobehavioral reviews, 52, 233–257. https://doi.org/10.1016/j.neubiorev.2015.03.002

Fernandez, T. V., & Leckman, J. F. (2016). Prenatal and Perinatal Risk Factors and the Promise of Birth Cohort Studies: Origins of Obsessive-Compulsive Disorder. JAMA psychiatry, 73(11), 1117–1118. https://doi.org/10.1001/jamapsychiatry.2016.2092

Fox, J., & Weisberg, S. (2018). car: Companion to Applied Regression. [R package]. Retrieved from https://cran.r-project.org/package=car.

Gauggel, S., Rieger, M., & Feghoff, T. A. (2004). Inhibition of ongoing responses in patients with Parkinson’s disease. Journal of neurology, neurosurgery, and psychiatry, 75(4), 539–544. https://doi.org/10.1136/jnnp.2003.016469

Gerdelat-Mas, A., Loubinoux, I., Tombari, D., Rascol, O., Chollet, F., & Simonetta-Moreau, M. (2005). Chronic administration of selective serotonin reuptake inhibitor (SSRI) paroxetine modulates human motor cortex excitability in healthy subjects. NeuroImage, 27(2), 314–322. https://doi.org/10.1016/j.neuroimage.2005.05.009

Grados M.A., Labuda M.C., Riddle M.A. & Walkup J.T. (1997). Obsessive-compulsive disorder in children and adolescents. International Review of Psychiatry, 9, 83–98. https://doi.org/10.1080/09540269775619

Guo, Y., Schmitz, T. W., Mur, M., Ferreira, C. S., & Anderson, M. C. (2018). A supramodal role of the basal ganglia in memory and motor inhibition: Meta-analytic evidence. Neuropsychologia, 108, 117–134. https://doi.org/10.1016/j.neuropsychologia.2017.11.033

Harville, D. A. (1974). Bayesian Inference for Variance Components Using Only Error Contrasts. Biometrika, 61(2), 383–385. https://doi.org/10.1093/biomet/61.2.383

Helbing, M.-L. C., & Ficca, M. (2009). Obsessive-Compulsive Disorder in School-Age Children. The Journal of School Nursing, 25(1), 15–26. https://doi.org/10.1177/1059840508328199

Higgins, J. P. T., & Thompson, S. G. (2002). Quantifying heterogeneity in a meta-analysis. Statistics in Medicine, 21(11), 1539–1558. https://doi.org/10.1002/sim.1186

Hiraoka, K., Kinoshita, A., Kunimura, H., & Matsuoka, M. (2018). Effect of variability of sequence length of go trials preceding a stop trial on ability of response inhibition in stop-signal task. Somatosensory & motor research, 35(2), 95–102. https://doi.org/10.1080/08990220.2018.1475351

Hsieh, S., & Lin, Y. C. (2017). Stopping ability in younger and older adults: Behavioral and event-related potential. Cognitive, affective & behavioral neuroscience, 17(2), 348–363. https://doi.org/10.3758/s13415-016-0483-7

Huedo-Medina, T. B., Sánchez-Meca, J., Marín-Martínez, F., & Botella, J. (2006). Assessing heterogeneity in meta-analysis: Q statistic or I2 index?. Psychological methods, 11(2), 193–206. https://doi.org/10.1037/1082-989X.11.2.193

Huizenga, H. M., van Bers, B. M., Plat, J., van den Wildenberg, W. P., & van der Molen, M. W. (2009). Task complexity enhances response inhibition deficits in childhood and adolescent attention-deficit/hyperactivity disorder: a meta-regression analysis. Biological psychiatry, 65(1), 39–45. https://doi.org/10.1016/j.biopsych.2008.06.021

Kalanthroff, E., Teichert, T., Wheaton, M. G., Kimeldorf, M. B., Linkovski, O., Ahmari, S. E., Fyer, A. J., Schneier, F. R., Anholt, G. E., & Simpson, H. B. (2017). The Role of Response Inhibition in Medicated and Unmedicated Obsessive-Compulsive Disorder Patients: Evidence from the Stop-Signal Task. Depression and anxiety, 34(3), 301–306. https://doi.org/10.1002/da.22492

Kessler, R. C., Petukhova, M., Sampson, N. A., Zaslavsky, A. M., & Wittchen, H. (2012). Twelve-month and lifetime prevalence and lifetime morbid risk of anxiety and mood disorders in the United States. International journal of methods in psychiatric research, 21(3), 169–184. https://doi.org/10.1002/mpr.1359

Lakens D. (2017). Equivalence Tests: A Practical Primer for *t* Tests, Correlations, and Meta-Analyses. Social psychological and personality science, 8(4), 355–362. https://doi.org/10.1177/1948550617697177

Li T, Higgins JPT, Deeks JJ (editors). Chapter 5: Collecting data. In: Higgins JPT, Thomas J, Chandler J, Cumpston M, Li T, Page MJ, Welch VA (editors). Cochrane Handbook for Systematic Reviews of Interventions version 6.1 (updated September 2020). Cochrane, 2020. Available from www.training.cochrane.org/handbook.

Liberati, A., Altman, D. G., Tetzlaff, J., Mulrow, C., Gøtzsche, P. C., Ioannidis, J. P., Clarke, M., Devereaux, P. J., Kleijnen, J., & Moher, D. (2009). The PRISMA statement for reporting systematic reviews and meta-analyses of studies that evaluate health care interventions: explanation and elaboration. Journal of clinical epidemiology, 62(10), e1–e34. https://doi.org/10.1016/j.jclinepi.2009.06.006

Lipszyc, J., & Schachar, R. (2010). Inhibitory control and psychopathology: a meta-analysis of studies using the stop signal task. Journal of the International Neuropsychological Society : JINS, 16(6), 1064–1076. https://doi.org/10.1017/S1355617710000895

Logan, G. D., & Cowan, W. B. (1984). On the ability to inhibit thought and action: A theory of an act of control. Psychological Review, 91(3), 295–327. https://doi.org/10.1037/0033-295X.91.3.295

Logan, G. D., Cowan, W. B., & Davis, K. A. (1984). On the ability to inhibit simple and choice reaction time responses: A model and a method. Journal of Experimental Psychology: Human Perception and Performance, 10(2), 276–291. https://doi.org/10.1037/0096-1523.10.2.276

Mattina, G. F., & Steiner, M. (2016). The need for inclusion of sex and age of onset variables in genetic association studies of obsessive-compulsive disorder: Overview. Progress in neuro-psychopharmacology & biological psychiatry, 67, 107–116. https://doi.org/10.1016/j.pnpbp.2016.01.012

Moher, D., Liberati, A., Tetzlaff, J., Altman, D. G., & PRISMA Group (2009). Preferred reporting items for systematic reviews and meta-analyses: the PRISMA statement. BMJ (Clinical research ed.), 339, b2535. https://doi.org/10.1136/bmj.b2535

Monzani, B., Rijsdijk, F., Harris, J., & Mataix-Cols, D. (2014). The structure of genetic and environmental risk factors for dimensional representations of DSM-5 obsessivecompulsive spectrum disorders. JAMA psychiatry, 71(2), 182–189. https://doi.org/10.1001/jamapsychiatry.2013.3524

Norman, L. J., Taylor, S. F., Liu, Y., Radua, J., Chye, Y., Wit, S. J. D., Huyser, D., Karahanoglu, F. I., Luks, T., Monarch, D., Mathews, C., Rubia, K., Suo, C., van den Heuvel, O. A., Yücel, M., Fitzgerald, K. (2019). Error Processing and Inhibitory Control in Obsessive-Compulsive Disorder: A Meta-analysis Using Statistical Parametric Maps. Biological Psychiatry, 85(9), 713–725. https://doi.org/10.1016/j.biopsych.2018.11.010

Ornstein, T. J., Levin, H. S., Chen, S., Hanten, G., Ewing-Cobbs, L., Dennis, M., Barnes, M., Max, J. E., Logan, G. D., & Schachar, R. (2009). Performance monitoring in children following traumatic brain injury. Journal of child psychology and psychiatry, and allied disciplines, 50(4), 506–513. https://doi.org/10.1111/j.1469-7610.2008.01997.x

Patterson, H., & Thompson, R. (1971). Recovery of Inter-Block Information when Block Sizes are Unequal. Biometrika, 58(3), 545–554. doi:10.2307/2334389

Penadés, R., Catalán, R., Rubia, K., Andrés, S., Salamero, M., & Gastó, C. (2007). Impaired response inhibition in obsessive compulsive disorder. European psychiatry : the journal of the Association of European Psychiatrists, 22(6), 404–410. https://doi.org/10.1016/j.eurpsy.2006.05.001

Pitman, R. K. (1987). A cybernetic model of obsessive-compulsive psychopathology. Comprehensive Psychiatry, 28(4), 334–343. https://doi.org/10.1016/0010-440x(87)90070-8

Pliszka, S. R., Liotti, M., Bailey, B. Y., Perez, R., 3rd, Glahn, D., & Semrud-Clikeman, M. (2007). Electrophysiological effects of stimulant treatment on inhibitory control in children with attention-deficit/hyperactivity disorder. Journal of child and adolescent psychopharmacology, 17(3), 356–366. https://doi.org/10.1089/cap.2006.0081

R Core Team. (2019). R: A Language and Environment for Statistical Computing. Vienna, Austria. Retrieved from https://www.R-project.org/

Robbins, T. W., Vaghi, M. M., & Banca, P. (2019). Obsessive-Compulsive Disorder: Puzzles and Prospects. Neuron, 102(1), 27–47. https://doi.org/10.1016/j.neuron.2019.01.046

Ruscio, A. M., Stein, D. J., Chiu, W. T., & Kessler, R. C. (2010). The epidemiology of obsessive-compulsive disorder in the National Comorbidity Survey Replication. Molecular psychiatry, 15(1), 53–63. https://doi.org/10.1038/mp.2008.94

Samuels J. & Nestadt G. (1997). Epidemiology and genetics of obsessive-compulsive disorder. International Review of Psychiatry, 9, 61–72. https://doi.org/10.1080/09540269775592

Schachar, R., & Logan, G. D. (1990). Impulsivity and inhibitory control in normal development and childhood psychopathology. Developmental Psychology, 26(5), 710–720. http://doi.org/10.1037/0012-1649.26.5.710

Sinopoli, K. J., Schachar, R., & Dennis, M. (2011). Reward improves cancellation and restraint inhibition across childhood and adolescence. Developmental psychology, 47(5), 1479–1489. https://doi.org/10.1037/a0024440

Skandali, N., Rowe, J. B., Voon, V., Deakin, J. B., Cardinal, R. N., Cormack, F., Passamonti, L., Bevan-Jones, W. R., Regenthal, R., Chamberlain, S. R., Robbins, T. W., & Sahakian, B. J. (2018). Dissociable effects of acute SSRI (escitalopram) on executive, learning and emotional functions in healthy humans. Neuropsychopharmacology : official publication of the American College of Neuropsychopharmacology, 43(13), 2645–2651. https://doi.org/10.1038/s41386-018-0229-z

Slusarek, M., Velling, S., Bunk, D., & Eggers, C. (2001). Motivational effects on inhibitory control in children with ADHD. Journal of the American Academy of Child and Adolescent Psychiatry, 40(3), 355–363. https://doi.org/10.1097/00004583-200103000-00016

Smith, J. L., Mattick, R. P., Jamadar, S. D., & Iredale, J. M. (2014). Deficits in behavioural inhibition in substance abuse and addiction: a meta-analysis. Drug and alcohol dependence, 145, 1–33. https://doi-org.myaccess.library.utoronto.ca/10.1016/j.drugalcdep.2014.08.009

Solanto, M. V., Abikoff, H., Sonuga-Barke, E., Schachar, R., Logan, G. D., Wigal, T., Hechtman, L., Hinshaw, S., & Turkel, E. (2001). The ecological validity of delay aversion and response inhibition as measures of impulsivity in AD/HD: a supplement to the NIMH multimodal treatment study of AD/HD. Journal of abnormal child psychology, 29(3), 215–228. https://doi.org/10.1023/a:1010329714819

Sterne, J. A., Sutton, A. J., Ioannidis, J. P., Terrin, N., Jones, D. R., Lau, J., Carpenter, J., Rücker, G., Harbord, R. M., Schmid, C. H., Tetzlaff, J., Deeks, J. J., Peters, J., Macaskill, P., Schwarzer, G., Duval, S., Altman, D. G., Moher, D., & Higgins, J. P. (2011). Recommendations for examining and interpreting funnel plot asymmetry in meta-analyses of randomised controlled trials. BMJ (Clinical research ed.), 343, d4002. https://doi.org/10.1136/bmj.d4002

Takeshima, N., Sozu, T., Tajika, A., Ogawa, Y., Hayasaka, Y., & Furukawa, T. A. (2014). Which is more generalizable, powerful and interpretable in meta-analyses, mean difference or standardized mean difference?. BMC medical research methodology, 14, 30. https://doi.org/10.1186/1471-2288-14-30

Tannock, R., Martinussen, R., & Frijters, J. (2000). Naming speed performance and stimulant effects indicate effortful, semantic processing deficits in attention-deficit/hyperactivity disorder. Journal of abnormal child psychology, 28(3), 237–252. https://doi.org/10.1023/a:1005192220001

Taylor S. (2011). Early versus late onset obsessive-compulsive disorder: evidence for distinct subtypes. Clinical psychology review, 31(7), 1083–1100. https://doi.org/10.1016/j.cpr.2011.06.007

Thakkar, K. N., Schall, J. D., Boucher, L., Logan, G. D., & Park, S. (2011). Response inhibition and response monitoring in a saccadic countermanding task in schizophrenia. Biological psychiatry, 69(1), 55–62. https://doi.org/10.1016/j.biopsych.2010.08.016

The jamovi project (2020). *jamovi*. (Version 1.2) [Computer Software]. Retrieved from https://www.jamovi.org.

van der Plas, E., Schachar, R. J., Hitzler, J., Crosbie, J., Guger, S. L., Spiegler, B. J., Ito, S., & Nieman, B. J. (2016). Brain structure, working memory and response inhibition in childhood leukemia survivors. Brain and behavior, 7(2), e00621. https://doi.org/10.1002/brb3.621

van Velzen, L. S., Vriend, C., de Wit, S. J., & van den Heuvel, O. A. (2014). Response inhibition and interference control in obsessive-compulsive spectrum disorders. Frontiers in human neuroscience, 8, 419. https://doi.org/10.3389/fnhum.2014.00419

Viechtbauer, W. (2010). Conducting Meta-Analyses in R with the metafor Package. Journal of Statistical Software, 36(3), 1–48. https://doi.org/10.18637/jss.v036.i03

Wells, G., Shea, B., O’Connell, D., Peterson, J., Welch, V., Losos, M., & Tugwell, P. (2019). The Newcastle-Ottawa Scale (NOS) for assessing the quality of nonrandomised studies in meta-analyses. Available from URL: http://www.ohri.ca/programs/clinical_epidemiology/oxford.asp (accessed May 2020).

