## Supplemental Information for "Obsessive Compulsive Disorder and Response Inhibition: Meta-analysis of the Stop-Signal Task"

S1. Literature Search Strategies.

S2. References of studies included in meta-analyses.
S3. Risk-of-bias evaluation for the included studies.
S4. PRISMA Checklist.
S5. Risk of bias funnel plot for SSRT and MRT in raw mean difference
S6. Forest Plot of Included Literature for SSRT and MRT in standardized mean difference
S7. Risk of Bias Funnel Plot for SSRT and MRT in standardized mean difference

***S1****.*

*Literature Search Strategies*

| Database [Platform] *Searches run November 28-29, 2020* | Results |
| --- | --- |
| Ovid MEDLINE: Epub Ahead of Print, In-Process & Other Non-Indexed Citations, Ovid MEDLINE® Daily and Ovid MEDLINE® 1946 – November 28 | 54 |
| Embase Classic+Embase 1947 to 2020 November 25 | 88 |
| APA PsycInfo 1806 to November Week 3 2020 | 45 |
| Web of Science [Clarivate Analytics] Nov 29, 2020 | 112 |
| TOTAL | 299 |

Ovid MEDLINE: Epub Ahead of Print, In-Process & Other Non-Indexed Citations, Ovid MEDLINE® Daily and Ovid MEDLINE® 1946 – November 28

Search Strategy:

| # | Searches | Results |
| --- | --- | --- |
| 1 | stop signal?.tw,kw. | 2301 |
| 2 | stop* task*.tw,kf. | 151 |
| 3 | (stop adj3 task*).tw,kw. | 1408 |
| 4 | or/1-3 | 2514 |
| 5 | Obsessive-Compulsive Disorder/ | 14564 |
| 6 | (anankastic personalit* or obsessive compulsive disorder? or obsessive compulsive neuros#s or OCD).tw,kw. | 15764 |
| 7 | or/5-6 | 20929 |
| 8 | "inhibition (psychology)"/ or proactive inhibition/ or reactive inhibition/ | 11365 |
| 9 | (inhibit* adj3 (respons* or behavi* or proactive* or reactive*)).tw,kw. | 61572 |
| 10 | (impulse? adj2 control*).tw,kw. | 3509 |
| 11 | or/8-10 | 73188 |
| 12 | executive function/ | 15222 |
| 13 | (executive adj2 (control or function*)).tw,kw. | 29872 |
| 14 | (cognitive adj2 (control or function*)).tw,kw. | 75968 |
| 15 | or/12-14 | 101825 |
| 16 | 4 and 7 | 65 |
| 17 | 11 or 15 | 170823 |
| 18 | 16 AND 17 | 54 |

Embase Classic+Embase 1947 to 2020 November 28

Search Strategy:

| # | Searches | Results |
| --- | --- | --- |
| 1 | stop signal?.tw,kw. | 2953 |
| 2 | stop task?.tw,kw. | 216 |
| 3 | (stop adj3 task*).tw,kw. | 1928 |
| 4 | or/1-3 | 3235 |
| 5 | obsessive compulsive disorder/ | 25862 |
| 6 | (anankastic personalit* or obsessive compulsive disorder? or obsessive compulsive neuros#s or OCD).tw,kw. | 23189 |
| 7 | or/5-6 | 34193 |
| 8 | "inhibition (psychology)"/ or proactive inhibition/ or reactive inhibition/ | 7931 |
| 9 | (inhibit* adj3 (respons* or behavi* or proactive* or reactive*)).tw,kw. | 79435 |
| 10 | (impulse? adj2 control*).tw,kw. | 5900 |
| 11 | or/8-10 | 90985 |
| 12 | Executive function/ | 42082 |
| 13 | (executive adj2 (control or function*)).tw,kw. | 46940 |
| 14 | (cognitive adj2 (control or function*)).tw,kw. | 114046 |
| 15 | Or/12-14 | 155539 |
| 16 | 11 or 15 | 241289 |
| 17 | 4 and 7 | 114 |
| 18 | 16 and 17 | 88 |

APA PsycInfo 1806 to November Week 3 2020

Search Strategy:

| # | Searches | Results |
| --- | --- | --- |
| 1 | stop signal?.tw. | 1659 |
| 2 | stop task?.tw. | 160 |
| 3 | (stop adj task*).tw. | 1328 |
| 4 | or/1-3 | 1823 |
| 5 | obsessive compulsive disorder/ | 14382 |
| 6 | (anankastic personalit* or obsessive compulsive disorder? or obsessive compulsive neuros#s or OCD).tw. | 17761 |
| 7 | or/5-6 | 19345 |
| 8 | Proactive inhibition/ or reactive inhibition/ or response inhibition/ or behavioral inhibition system/ | 4233 |
| 9 | (inhibit* adj3 (respons* or behavi* or proactive* or reactive*)).tw. | 16247 |
| 10 | (impulse? adj2 control*).tw. | 4824 |
| 11 | or/8-10 | 22514 |
| 12 | Executive function/ | 10774 |
| 13 | (executive adj2 (control or function*)).tw. | 31763 |
| 14 | (cognitive adj2 (control or function*)).tw. | 56113 |
| 15 | or/12-14 | 80952 |
| 16 | 11 or 15 | 100383 |
| 17 | 4 and 7 | 62 |
| 18 | 16 and 17 | 45 |

Web of Science [v.5.35] Nov 28, 2020

Search Strategy:

| Set | Results | Search |
| --- | --- | --- |
| #14 | 112 | #12 AND #13  Indexes=SCI-EXPANDED, SSCI, A&HCI, CPCI-S, CPCI-SSH, BKCI-S, BKCI-SSH, ESCI Timespan=All years |
| #13 | 227,943 | #8 OR #11  Indexes=SCI-EXPANDED, SSCI, A&HCI, CPCI-S, CPCI-SSH, BKCI-S, BKCI-SSH, ESCI Timespan=All years |
| #12 | 130 | #4 AND #5  Indexes=SCI-EXPANDED, SSCI, A&HCI, CPCI-S, CPCI-SSH, BKCI-S, BKCI-SSH, ESCI Timespan=All years |
| #11 | 141,756 | #9 OR #10  Indexes=SCI-EXPANDED, SSCI, A&HCI, CPCI-S, CPCI-SSH, BKCI-S, BKCI-SSH, ESCI Timespan=All years |
| #10 | 106,848 | TS=(cognitive NEAR/2 (control or function) )  Indexes=SCI-EXPANDED, SSCI, A&HCI, CPCI-S, CPCI-SSH, BKCI-S, BKCI-SSH, ESCI Timespan=All years |
| #9 | 49,437 | TS=(executive NEAR/2 (control or function) )  Indexes=SCI-EXPANDED, SSCI, A&HCI, CPCI-S, CPCI-SSH, BKCI-S, BKCI-SSH, ESCI Timespan=All years |
| #8 | 92,299 | #6 OR #7  Indexes=SCI-EXPANDED, SSCI, A&HCI, CPCI-S, CPCI-SSH, BKCI-S, BKCI-SSH, ESCI Timespan=All years |
| #7 | 6,770 | TS=((impulse* NEAR/2 control*))  Indexes=SCI-EXPANDED, SSCI, A&HCI, CPCI-S, CPCI-SSH, BKCI-S, BKCI-SSH, ESCI Timespan=All years |
| #6 | 85,988 | TS=((inhibit* NEAR/3 (respons* or behavi* or proactive* or reactive*) ))  Indexes=SCI-EXPANDED, SSCI, A&HCI, CPCI-S, CPCI-SSH, BKCI-S, BKCI-SSH, ESCI Timespan=All years |
| #5 | 26,710 | TS=(("anankastic personalit*" or "obsessive compulsive disorder*" or "obsessive compulsive neurosis" or "obsessive compulsive neuroses" or OCD))  Indexes=SCI-EXPANDED, SSCI, A&HCI, CPCI-S, CPCI-SSH, BKCI-S, BKCI-SSH, ESCI Timespan=All years |
| #4 | [3,525](https://apps-webofknowledge-com.myaccess.library.utoronto.ca/summary.do?product=WOS&doc=1&qid=7&SID=6DX49XyX8z1jNBNC1Dk&search_mode=AdvancedSearch&update_back2search_link_param=yes) | #1 OR #2 OR #3  Indexes=SCI-EXPANDED, SSCI, A&HCI, CPCI-S, CPCI-SSH, BKCI-S, BKCI-SSH, ESCI Timespan=All years |
| #3 | 1,889 | TS=(stop NEAR/2 task*)  Indexes=SCI-EXPANDED, SSCI, A&HCI, CPCI-S, CPCI-SSH, BKCI-S, BKCI-SSH, ESCI Timespan=All years |
| #2 | 192 | TS=("stop task*")  Indexes=SCI-EXPANDED, SSCI, A&HCI, CPCI-S, CPCI-SSH, BKCI-S, BKCI-SSH, ESCI Timespan=All years |

***S2****.*

*References of Studies Included in Meta-Analyses*

| Author (year) | Reference |
| --- | --- |
| Bersani (2013) | Bersani, G., Quartini, A., Ratti, F., Pagliuca, G., & Gallo, A. (2013). Olfactory identification deficits and associated response inhibition in obsessive-compulsive disorder: On the scent of the orbitofronto-striatal model. *Psychiatry Research*, *210*(1), 208–214.<https://doi.org/10.1016/j.psychres.2013.05.032> |
| Boisseau (2012) | Boisseau, C. L., Thompson-Brenner, H., Caldwell-Harris, C., Pratt, E., Farchione, T., & Harrison Barlow, D. (2012). Behavioral and cognitive impulsivity in obsessive-compulsive disorder and eating disorders. *Psychiatry Research*, *200*(2–3), 1062–1066.<https://doi.org/10.1016/j.psychres.2012.06.010> |
| Chamberlain (2006) | Chamberlain, S. R., Fineberg, N. A., Blackwell, A. D., Clark, L., Robbins, T. W., & Sahakian, B. J. (2006). A neuropsychological comparison of obsessive-compulsive disorder and trichotillomania. *Neuropsychologia*, *45*, 654–662.<https://doi.org/10.1016/j.neuropsychologia.2006.07.016> |
| Chamberlain (2007) | Chamberlain, S. R., Naomi Fineberg, M. A., Lara Menzies, M. A., Andrew Blackwell, B. D., Bullmore, E. T., Trevor Robbins, Bc. W., & Sahakian, B. J. (2007). Impaired Cognitive Flexibility and Motor Inhibition in Unaffected First-Degree Relatives of Patients With Obsessive-Compulsive Disorder. In *Am J Psychiatry* (Vol. 164). |
| De Wit (2012) | De Wit, S. J., De Vries, F. E., Van Der Werf, Y. D., Cath, D. C., Heslenfeld, D. J., Veltman, E. M., Van Balkom, A. J. L. M., Veltman, D. J., & Van Den Heuvel, O. A. (2012). Presupplementary motor area hyperactivity during response inhibition: A candidate endophenotype of obsessive-compulsive disorder. *American Journal of Psychiatry*, *169*(10), 1100–1108.<https://doi.org/10.1176/appi.ajp.2012.12010073> |
| Fan (2016) | Fan, J., Liu, W., Lei, H., Cai, L., Zhong, M., Dong, J., Zhou, C., & Zhu, X. (2016). Components of inhibition in autogenous- and reactive-type obsessive-compulsive disorder: Dissociation of interference control. *Biological Psychology*, *117*, 117–130.<https://doi.org/10.1016/j.biopsycho.2016.03.008> |
| Frydman (2020) | Frydman, I., Mattos, P., de Oliveira-Souza, R., Yücel, M., Chamberlain, S. R., Moll, J., & Fontenelle, L. F. (2020). Self-reported and neurocognitive impulsivity in obsessive-compulsive disorder. *Comprehensive Psychiatry*, *97*.<https://doi.org/10.1016/j.comppsych.2019.152155> |
| Gooskens (2018) | Gooskens, B., Bos, D. J., Mensen, V. T., Shook, D. A., Bruchhage, M. M. K., Naaijen, J., Wolf, I., Brandeis, D., Williams, S. C. R., Buitelaar, J. K., Oranje, B., & Durston, S. (2019). No evidence of differences in cognitive control in children with autism spectrum disorder or obsessive-compulsive disorder: An fMRI study. *Developmental Cognitive Neuroscience*, *36*.<https://doi.org/10.1016/j.dcn.2018.11.004> |
| Hampshire (2019) | Hampshire, A., Zadel, A., Sandrone, S., Soreq, E., Fineberg, N., Bullmore, E. T., Robbins, T. W., Sahakian, B. J., & Chamberlain, S. R. (2020). Inhibition-Related Cortical Hypoconnectivity as a Candidate Vulnerability Marker for Obsessive-Compulsive Disorder. *Biological Psychiatry: Cognitive Neuroscience and Neuroimaging*, *5*(2), 222–230.<https://doi.org/10.1016/j.bpsc.2019.09.010> |
| Heinzel (2018) | Heinzel, S., Kaufmann, C., Grützmann, R., Hummel, R., Klawohn, J., Riesel, A., Bey, K., Lennertz, L., Wagner, M., & Kathmann, N. (2018). Neural correlates of working memory deficits and associations to response inhibition in obsessive compulsive disorder. *NeuroImage: Clinical*, *17*, 426–434.<https://doi.org/10.1016/j.nicl.2017.10.039> |
| Kang (2013) | Kang, D. H., Jang, J. H., Han, J. Y., Kim, J. H., Jung, W. H., Choi, J. S., Choi, C. H., & Kwon, J. S. (2013). Neural correlates of altered response inhibition and dysfunctional connectivity at rest in obsessive-compulsive disorder. *Progress in Neuro-Psychopharmacology and Biological Psychiatry*, *40*(1), 340–346.<https://doi.org/10.1016/j.pnpbp.2012.11.001> |
| Lei (2015) | Lei, H., Zhu, X., Fan, J., Dong, J., Zhou, C., Zhang, X., & Zhong, M. (2015). Is impaired response inhibition independent of symptom dimensions in obsessive-compulsive disorder? Evidence from ERPs. *Scientific Reports*, *5*.<https://doi.org/10.1038/srep10413> |
| Lei (2017) | Lei, H., Zhong, M., Fan, J., Zhang, X., Cai, L., & Zhu, X. (2017). Age at symptom onset is not associated with reduced action cancelation in adults with obsessive-compulsive disorder. *Psychiatry Research*, *252*, 180–184.<https://doi.org/10.1016/j.psychres.2017.02.063> |
| Negreiros (2020) | Negreiros, J., Belschner, L., Best, J. R., Lin, S., Franco Yamin, D., Joffres, Y., Selles, R. R., Jaspers-Fayer, F., Miller, L. D., Woodward, T. S., Honer, W. G., & Stewart, S. E. (2020). Neurocognitive risk markers in pediatric obsessive–compulsive disorder. *Journal of Child Psychology and Psychiatry and Allied Disciplines*, *61*(5), 605–613.<https://doi.org/10.1111/jcpp.13153> |
| Ornstein (2010) | Ornstein, T. J., Arnold, P., Manassis, K., Mendlowitz, S., & Schachar, R. (2010). Neuropsychological performance in childhood OCD: A preliminary study. *Depression and Anxiety*, *27*(4), 372–380.<https://doi.org/10.1002/da.20638> |
| Penad**é**s (2007) | Penadés, R., Catalán, R., Rubia, K., Andrés, S., Salamero, M., & Gastó, C. (2007). Impaired response inhibition in obsessive compulsive disorder. *European Psychiatry*, *22*(6), 404–410.<https://doi.org/10.1016/j.eurpsy.2006.05.001> |
| Sohn (2014) | Sohn, S. Y., Kang, J. I., Namkoong, K., & Kim, S. J. (2014). Multidimensional Measures of Impulsivity in Obsessive-Compulsive Disorder: Cannot Wait and Stop. *PLoS ONE*, *9*(11).<https://doi.org/10.1371/journal.pone.0111739> |
| Suñol (2019) | Suñol, M., Martínez-Zalacaín, I., Picó-Pérez, M., López-Solà, C., Real, E., Fullana, M. À., Pujol, J., Cardoner, N., Menchón, J. M., Alonso, P., & Soriano-Mas, C. (2020). Differential patterns of brain activation between hoarding disorder and obsessive-compulsive disorder during executive performance. *Psychological Medicine*, *50*(4), 666–673.<https://doi.org/10.1017/S0033291719000515> |
| Thorsen (2020) | Lillevik Thorsen, A., de Wit, S. J., Hagland, P., Ousdal, O. T., Hansen, B., Hagen, K., Kvale, G., & van den Heuvel, O. A. (2020). Stable inhibition-related inferior frontal hypoactivation and fronto-limbic hyperconnectivity in obsessive–compulsive disorder after concentrated exposure therapy. *NeuroImage: Clinical*, *28*, 102460. https://doi.org/https://doi.org/10.1016/j.nicl.2020.102460 |
| Woolley (2008) | Woolley, J., Heyman, I., Brammer, M., Frampton, I., McGuire, P. K., & Rubia, K. (2008). Brain activation in paediatric obsessive-compulsive disorder during tasks of inhibitory control. *British Journal of Psychiatry*, *192*(1), 25–31.<https://doi.org/10.1192/bjp.bp.107.036558> |
| Yu (2019) | Yu, F., Chen, X., Zhang, L., Bai, T., Gao, Y., Dong, Y., Luo, Y., Zhu, C., & Wang, K. (2019). Shared Response Inhibition Deficits but Distinct Error Processing Capacities Between Schizophrenia and Obsessive-Compulsive Disorder Patients Revealed by Event-Related Potentials and Oscillations During a Stop Signal Task. *Frontiers in Psychiatry*, *10*. https://doi.org/10.3389/fpsyt.2019.00853 |

*Note.* N = 21 included studies.

***S3****.*

*Risk-of-Bias Evaluation for the Included Studies*

| Author | Selection (maximum four stars) | Comparability (maximum two stars) | Outcome (maximum three stars) | Stars allocated to study/total potential stars |
| --- | --- | --- | --- | --- |
| Bersani (2013) | **** | ** | *** | 9/9 |
| Boisseau (2012) | ** | ** |  | 4/9 |
| Chamberlain (2006) | ** | ** | ** | 6/9 |
| Chamberlain (2007) | **** | ** | ** | 8/9 |
| De Wit (2012) | **** | ** | * | 8/9 |
| Fan (2016) | **** | ** | * | 7/9 |
| Frydman (2020) | **** | ** | *** | 9/9 |
| Gooskens (2018) | **** | ** | * | 7/9 |
| Hampshire (2019) | **** | ** | ** | 8/9 |
| Heinzel (2018) | **** | ** | * | 7/9 |
| Kang (2013) | *** | ** | ** | 7/9 |
| Lei (2015) | **** | ** | ** | 8/9 |
| Lei (2017) | *** | * | * | 5/9 |
| Negreiros (2020) | ** | * | * | 4/9 |
| Ornstein (2010) | **** | ** | * | 7/9 |
| Penades (2007) | ** | ** | ** | 6/9 |
| Sohn (2014) | *** | ** | ** | 7/9 |
| Sunol (2019) | **** | ** | *** | 9/9 |
| Thorsen (2019) | **** | ** | *** | 9/9 |
| Woolley (2008) | * | ** | *** | 6/9 |
| Yu (2019) | **** | ** | * | 7/9 |

*Note.* N = 21 included studies. * denotes a study having met the risk of bias assessment criteria. A lower number received by a study denotes a greater chance of bias in each respective category.

**S4**.

*PRISMA Checklist*

| Section/Topic | # | Checklist Item | Reported on page # |
| --- | --- | --- | --- |
| TITLE | | | |
| TItle | 1 | Identify the report as a systematic review, meta-analysis, or both. | 1 |
| ABSTRACT | | | |
| Structured summary | 2 | Provide a structured summary including, as applicable: background; objectives; data sources; study eligibility criteria, participants, and interventions; study appraisal and synthesis methods; results; limitations; conclusions and implications of key findings; systematic review registration number. | 3 |
| INTRODUCTION | | | |
| Rationale | 3 | Describe the rationale for the review in the context of what is already known. | 4-6 |
| Objectives | 4 | Provide an explicit statement of questions being addressed with reference to participants, interventions, comparisons, outcomes, and study design (PICOS). | 6 |
| METHODS | | | |
| Protocol and registration | 5 | Indicate if a review protocol exists, if and where it can be accessed (e.g., Web address), and, if available, provide registration information including registration number. | 6  PROSPERO (No. 212770) |
| Eligibility criteria | 6 | Specify study characteristics (e.g., PICOS, length of follow-up) and report characteristics (e.g., years considered, language, publication status) used as criteria for eligibility, giving rationale. | 7 |
| Information sources | 7 | Describe all information sources (e.g., databases with dates of coverage, contact with study authors to identify additional studies) in the search and date last searched. | 7 |
| Search | 8 | Present full electronic search strategy for at least one database, including any limits used, such that it could be repeated. | Supplemental 1 |
| Study selection | 9 | State the process for selecting studies (i.e., screening, eligibility, included in systematic review, and, if applicable, included in the meta-analysis). | 7-8 |
| Data collection process | 10 | Describe method of data extraction from reports (e.g., piloted forms, independently, in duplicate) and any processes for obtaining and confirming data from investigators. | 8 |
| Data items | 11 | List and define all variables for which data were sought (e.g., PICOS, funding sources) and any assumptions and simplifications made. | 8 |
| Risk of bias in individual studies | 12 | Describe methods used for assessing risk of bias of individual studies (including specification of whether this was done at the study or outcome level), and how this information is to be used in any data synthesis. | 8 |
| Summary measures | 13 | State the principal summary measures (e.g., risk ratio, difference in means). | 9 |
| Synthesis of results | 14 | Describe the methods of handling data and combining results of studies, if done, including measures of consistency (e.g., I^2^) for each meta-analysis. | 9-10 |
| Risk of bias across studies | 15 | Specify any assessment of risk of bias that may affect the cumulative evidence (e.g., publication bias, selective reporting within studies). | 10 |
| Additional analyses | 16 | Describe methods of additional analyses (e.g., sensitivity or subgroup analyses, meta-regression), if done, indicating which were pre-specified. | 9 |
| RESULTS | | | |
| Study selection | 17 | Give numbers of studies screened, assessed for eligibility, and included in the review, with reasons for exclusions at each stage, ideally with a flow diagram. | 10  Figure 1 |
| Study characteristics | 18 | For each study, present characteristics for which data were extracted (e.g., study size, PICOS, follow-up period) and provide the citations. | 10  Table 1  Supplemental 2 |
| Risk of bias within studies | 19 | Present data on risk of bias of each study and, if available, any outcome-level assessment (see Item 12). | 8  Supplemental 3 |
| Results of individual studies | 20 | For all outcomes considered (benefits or harms), present, for each study: (a) simple summary data for each intervention group and (b) effect estimates and confidence intervals, ideally with a forest plot. | 10  Figure 2 a)  Figure 2 b)  Supplemental 6 |
| Synthesis of results | 21 | Present results of each meta-analysis done, including confidence intervals and measures of consistency. | 11 |
| Risk of bias across studies | 22 | Present results of any assessment of risk of bias across studies (see item 15) | 12  Supplemental 5  Supplemental 7 |
| Additional analysis | 23 | Give results of additional analyses, if done (such as sensitivity or subgroup analyses, meta-regression) (see item 16) | 11 |
| DISCUSSION | | | |
| Summary of evidence | 23 | Summarise the main findings including the strength of evidence for each main outcome; consider their relevance to key groups (such as health care providers, users, and policy makers) | 12-13 |
| Limitations | 25 | Discuss limitations at study and outcome level (such as risk of bias), and at review level (such as incomplete retrieval of identified research, reporting bias | 14-16 |
| Conclusion | 26 | Provide a general interpretation of the results in the context of other evidence, and implications for future research | 17-18 |
| FUNDING | | | |
| Funding | 27 | Describe sources of funding for the systematic review and other support (such as supply of data) and role of funders for the systematic review | 2 |

Moher, D., Liberati, A., Tetzlaff, J., Altman, D. G., & PRISMA Group (2009). Preferred reporting items for systematic reviews and meta-analyses: the PRISMA statement. *BMJ (Clinical research ed.)*, *339*, b2535. <https://doi.org/10.1136/bmj.b2535>

***S5****.*

*Risk of Bias Funnel Plot for (a) SSRT and (b) MRT*

**Figure S5a**.

*Funnel Plot of Included Literature Concerning SSRT Scores Between OCD Patients and Healthy Controls in raw mean difference*
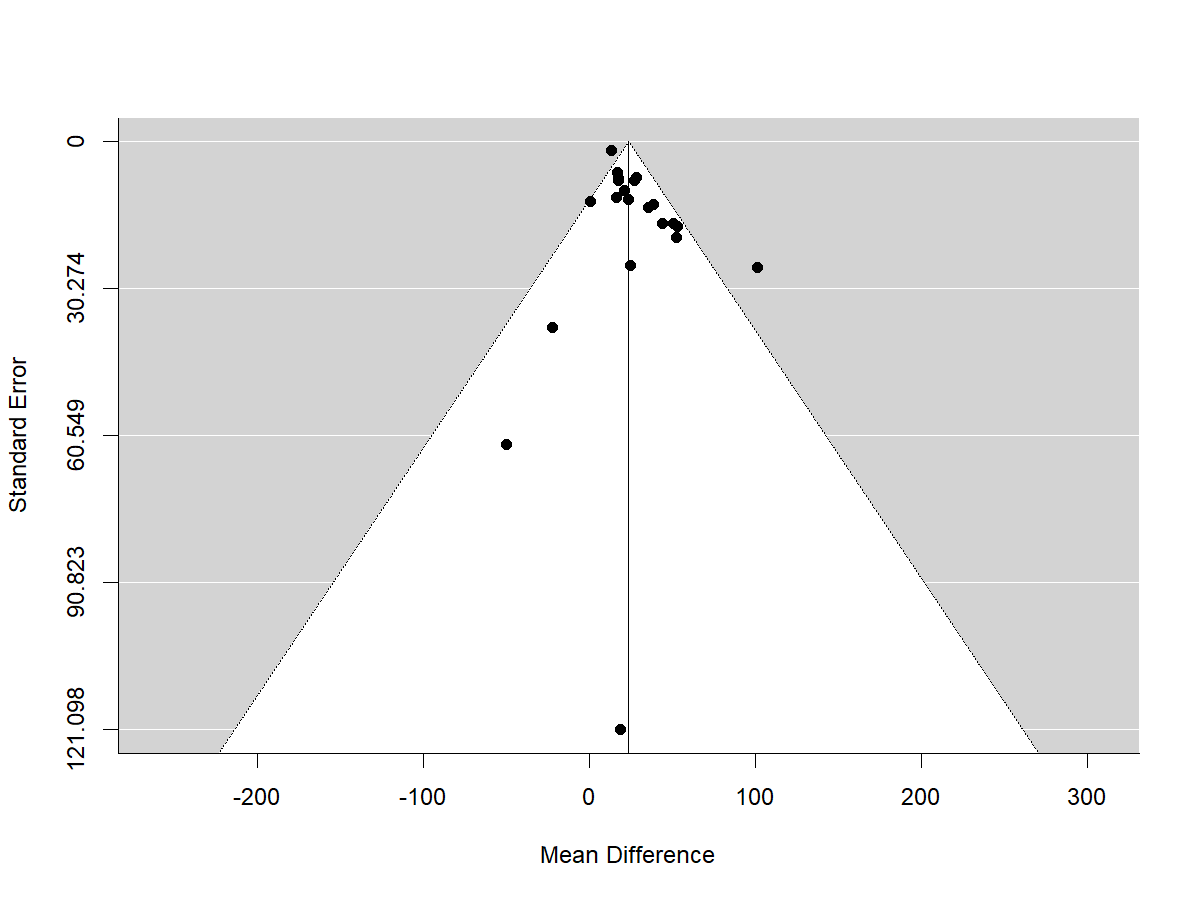

*Note.* N = 21 included studies. Funnel plot for the risk of bias across studies for the random-effects meta-analysis of SSRT scores. The mean difference was plotted against standard error for all studies that reported SSRT scores and revealed significant asymmetry (Z = 3.66; *p* < 0.001).

**Figure S5b**.

*Funnel Plot of Included Literature Concerning MRT Scores Between OCD Patients and Healthy*

*Controls in raw mean difference*

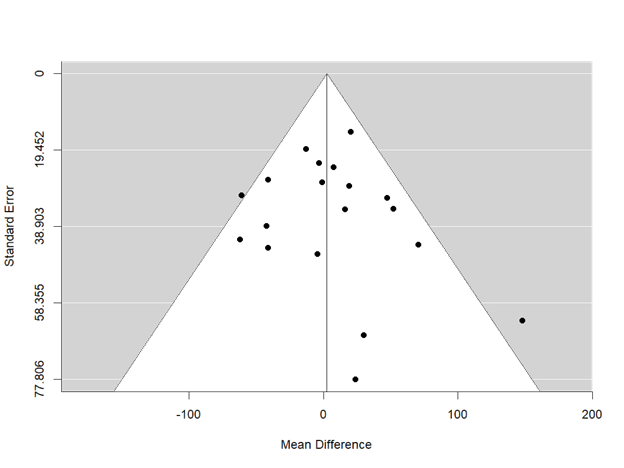

*Note.* N = 19 included studies. Funnel plot for the risk of bias across studies for the random-effects meta-analysis of MRT scores. The mean difference was plotted against standard error for all studies that reported MRT scores and revealed nonsignificant asymmetry (Z = 0.77; *p* = 0.439).

**Figure S5c.**

*Funnel Plot of Included Literature Concerning SSRT Scores Between OCD Patients and Healthy Controls with Outliers (Negreios et al., 2020 & Penades et al., 2007) in raw mean difference*

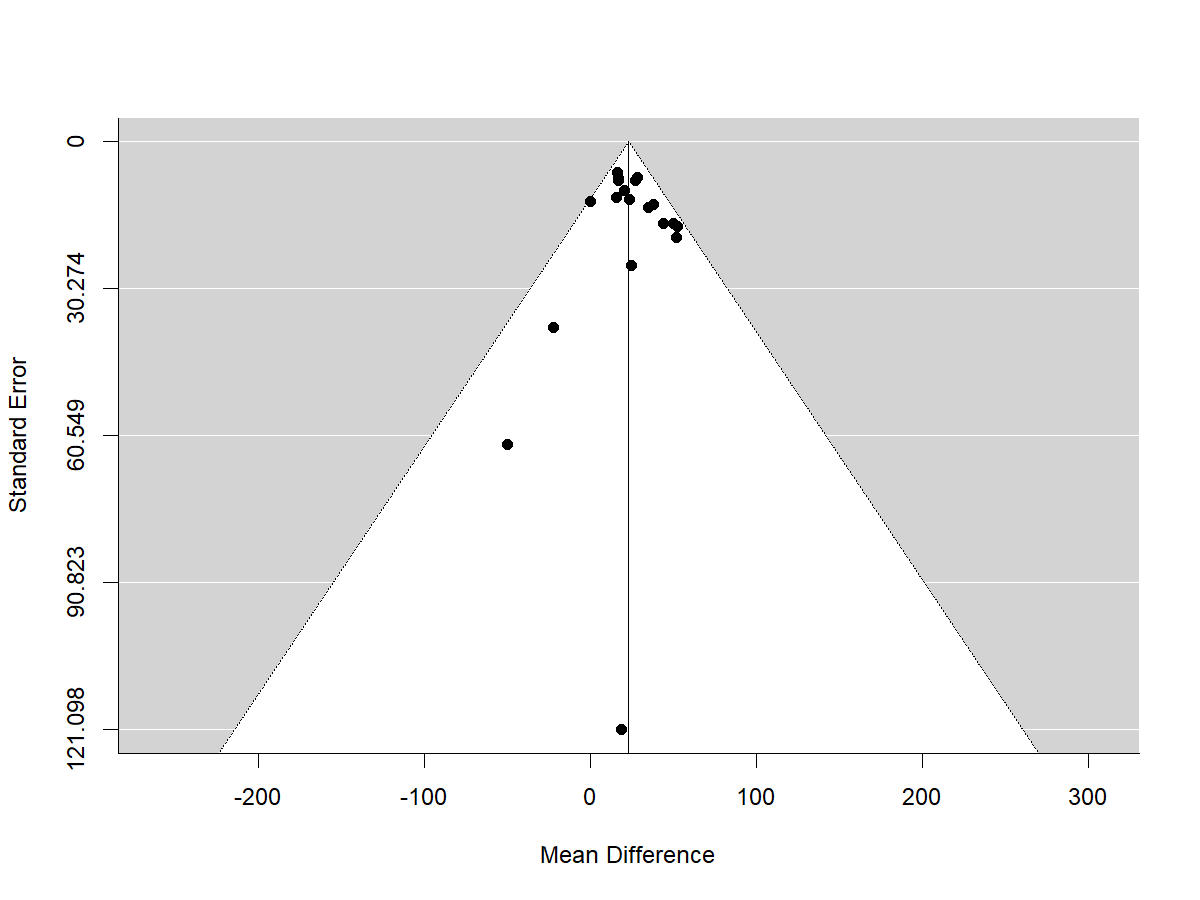
*Note.* Funnel plot for the risk of bias across studies for the random-effects meta-analysis of SSRT scores, with study outliers (Negreios et al., 2020 & Penades et al., 2007) removed. The mean difference was plotted against standard error for all studies that reported SSRT scores and revealed nonsignificant asymmetry (Z = 0.735, *p* = 0.462).

***S6.***

*Forest Plots of Included Literature for (a) SSRT and (b) MRT in standardized mean difference*

**Figure S6a**.

*Forest Plot of Included Literature Concerning SSRT Scores Between OCD Patients and Healthy Controls in standardized mean difference*

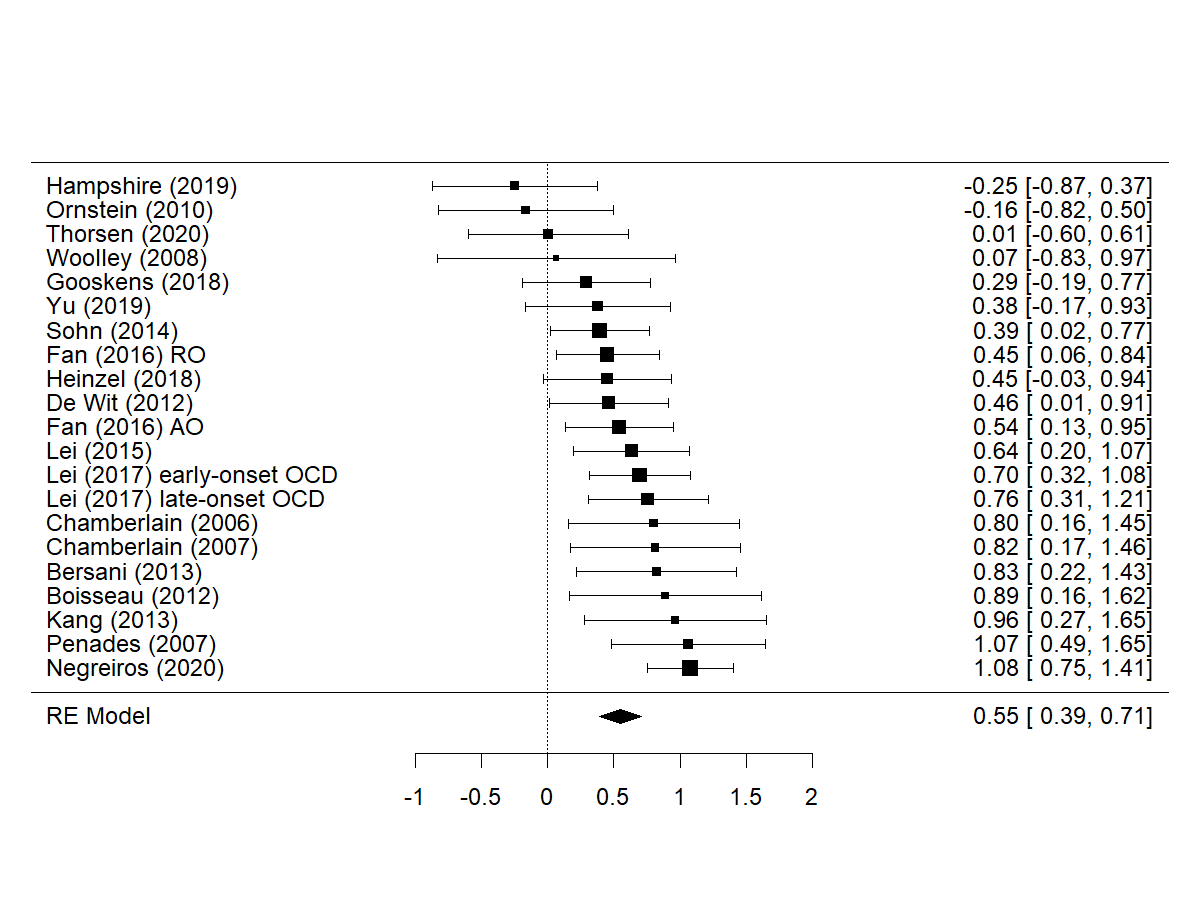

*Note.* Hedges’ g = 0.552 ; p <0.001 ; 95% CI [0.394, 0.710]; Heterogeneity: I^2^ = 50.24%; Q = 36.497; *p* = 0.013. This figure demonstrates the weighted random effects model of standardized mean difference scores using Hedge's g as the model estimator of SSRT (n= 21). RE = random effects, as estimated through restricted maximum likelihood. AO = autogenous obsessions. RO = reactive obsessions.

**Figure S6B**.

*Forest Plot of Included Literature Concerning MRT Scores Between OCD Patients and Healthy Controls in standardized mean difference*

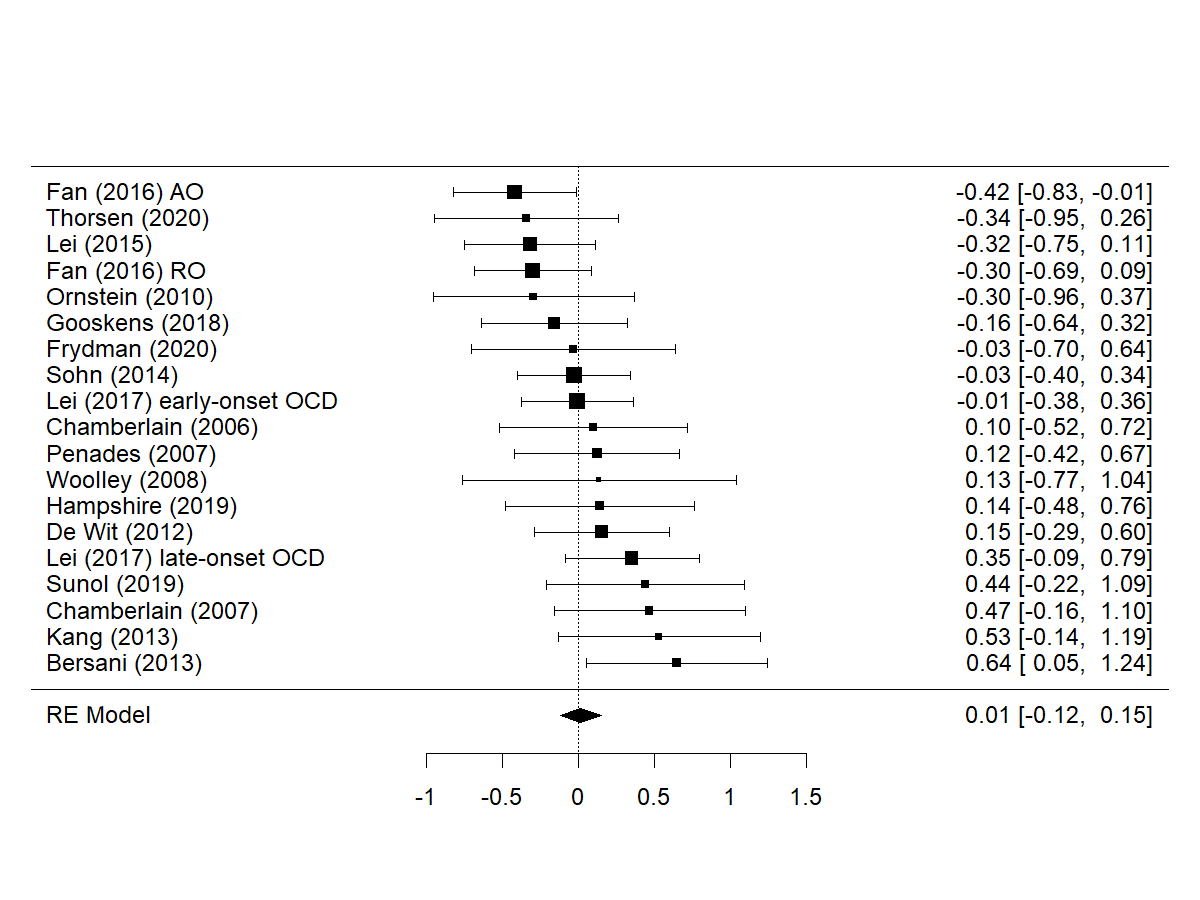

*Note.* Hedges’ g = 0.0144; p = 0.832; 95% CI [-0.119, 0.148]; Heterogeneity: I^2^ = 21.09%; Q = 25.214; *p* = 0.119. This figure demonstrates the weighted random effects model of the standardized mean difference scores using Hedge's g as the model estimator for studies that included MRT (n= 19). RE = random effects, as estimated through restricted maximum likelihood. AO = autogenous obsessions. RO = reactive obsessions.

***S7****.*

*Risk of Bias Funnel Plot for (a) SSRT and (b) MRT in standardized mean difference*

**Figure S7a**.

*Funnel Plot of Included Literature Concerning SSRT Scores Between OCD Patients and Healthy Controls in standardized mean difference*

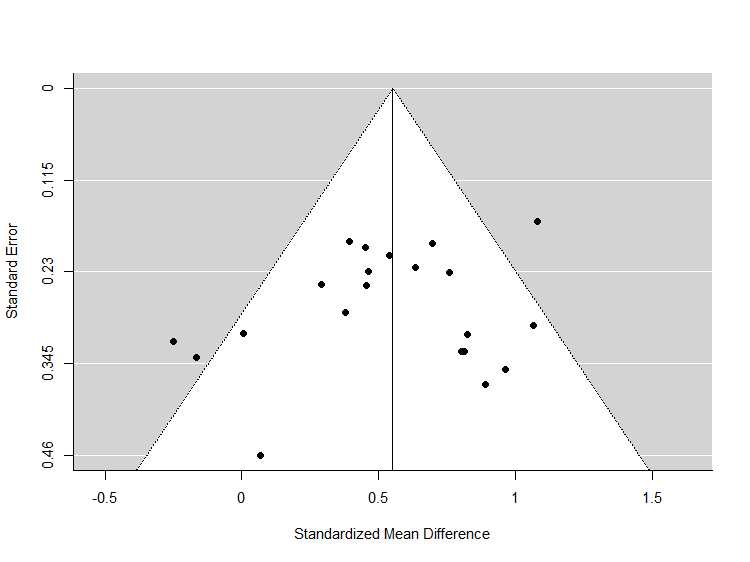

*Note.* N = 21 included studies. Funnel plot for the risk of bias across studies for the random-effects meta-analysis of SSRT scores. Hedge's g standardized mean difference was plotted against standard error for all studies that reported SSRT scores and revealed nonsignificant asymmetry (Z = -0.964; *p* = 0.335).

**Figure S7b**.

*Funnel Plot of Included Literature Concerning MRT Scores Between OCD Patients and Healthy*

*Controls in standardized mean difference*

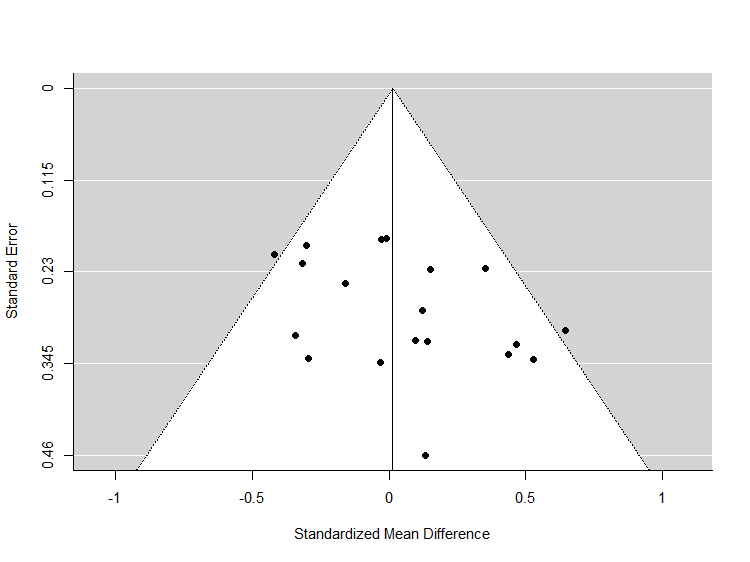

*Note.* N = 19 included studies. Funnel plot for the risk of bias across studies for the random-effects meta-analysis of MRT scores. Hedge's g standardized mean difference was plotted against standard error for all studies that reported MRT scores and revealed nonsignificant asymmetry (Z = 1.901; *p* = 0.057).
